# Transcriptomic Profiling of Bronchial Epithelium Reveals Dysregulated Interferon and Inflammatory Responses to Rhinovirus in Exacerbation-Prone Pediatric Asthma

**DOI:** 10.1101/2025.10.19.683298

**Authors:** Naresh Doni Jayavelu, Basilin Benson, Patricia C. dela Cruz, Weston T. Powell, Lucille M. Rich, Elizabeth R. Vanderwall, Camile R. Gates, Andrew J Nagel, Maria P. White, Nyssa B. Samanas, Kourtnie Whitfield, Teal S. Hallstrand, Steven F. Ziegler, Matthew C. Altman, Jason S. Debley

**Author notes:** **These authors contributed equally to this work as first authors** (Naresh Doni Jayavelu, Basilin Benson). **These authors contributed equally to this work as senior authors** (Matthew C. Altman, Jason S. Debley). **Corresponding Author:** Jason S. Debley, MD, MPH; ADDRESS: Center for Respiratory Biology and Therapeutics, Seattle Children’s Research Institute, 1916 Boren Ave., Seattle, WA, 98101.

## Abstract

Host factors influencing susceptibility to rhinovirus-induced asthma exacerbations remain poorly characterized. Using organotypic bronchial epithelial cultures from well-characterized children with asthma and healthy children, this study investigated viral load kinetics and resultant host responses by bulk and single-cell transcriptomics and targeted protein analyses. Bronchial epithelium from exacerbation-prone children exhibited greater rhinovirus replication and a cascade of exaggerated downstream interferon (IFN), inflammatory, epithelial stress, and remodeling responses. These transcriptional patterns were confirmed and further refined using single-cell transcriptomics, revealing cell type-specific contributions—particularly from non-ciliated cell populations including secretory immune response, tuft, and basal cells. We observed that these post-infection differences were associated with lower pre-infection IFN-stimulated gene (ISG) expression and protein levels of the ISG CXCL10. Prophylactic IFN-β treatment reduced viral replication and normalized downstream responses, supporting low baseline (pre-infection) IFN tone as a modifiable causal determinant of host susceptibility to adverse rhinovirus-induced responses in exacerbation-prone children with asthma.

## Introduction

Viral infections are the primary trigger of up to 90% of asthma exacerbations in children, with rhinoviruses (RV) being the most strongly associated with exacerbations (1, 2). Host factors influencing the risk of RV-triggered exacerbations remain poorly characterized. Early studies suggested that primary bronchial epithelial cells (BECs) from asthma donors had deficient type I and/or III interferon (IFN) responses to RV (3, 4) and postulated that deficient epithelial IFN responses to viruses may predispose to exacerbations. However, this concept remained controversial over the past two decades as later studies did not report differences in epithelial IFN responses to viruses between asthma and healthy donors (5, 6). More recently, we and others observed that, in fact, greater BEC IFN stimulated gene (ISG) expression *in vivo* (7) or in response to *ex vivo* viral infection (8) is associated with lower lung function in asthma donors. Furthermore, a recent case-control study of asthmatic children utilizing nasal lavage samples prospectively through viral upper respiratory tract infections, reported greater upregulation of ISG responses during viral illnesses that triggered acute exacerbations as compared to illnesses that did not, that this greater ISG expression associated with greater RV quantity and lower lung function throughout illness, and that lower pre-infection ISG expression associated with higher post-infection ISG expression and predicted higher risk for exacerbation (9, 10). These results suggested that low baseline airway interferon tone in children with asthma may permit accelerated viral replication and exaggerated host responses underpinning increased exacerbation risk, but this hypothesis remained to be tested.

To directly test this hypothesis, we leveraged BEC organotypic cultures derived from well-characterized pediatric donors with and without asthma. While nasal samples are practical for clinical studies, they are not directly sampling the lower airway, where asthma is manifest, and they represent mixtures of epithelial and immune cells characterizing a primary and secondary host response to virus. In contrast BEC cultures represent the lower airway and are pure epithelial cells that retain key structural and functional features of the host’s native airway epithelium, including mucociliary differentiation, innate immune signaling capacity, and donor-specific transcriptional programs. This model system enables precise temporal dissection of host-viral dynamics and allows for controlled perturbation of the pre-infection epithelial IFN milieu. By sampling from children with and without history of severe asthma exacerbations as well as healthy children without asthma, and experimentally modulating baseline IFN tone prior to RV infection, we aimed to determine whether low tonic IFN activity is a modifiable determinant of epithelial susceptibility to viral replication, downstream inflammatory responses, and exacerbation propensity. This approach offers a mechanistically grounded framework to identify epithelial immune phenotypes that predispose to exacerbation and may inform preventative or therapeutic strategies targeting epithelial antiviral readiness in children with asthma.

## Results

### Increased RV replication in BECs from children with severe asthma exacerbations

Primary BECs were collected from children with asthma (n=37) and healthy children (HC; n=3). Among children with asthma, 23 had a history of severe exacerbation (SE) and 14 children had no severe exacerbation history (NSE) (Table 1). Severe exacerbation history was defined as history of exacerbation(s) requiring systemic corticosteroids and/or emergency department care/hospitalization for asthma (11). All three groups had similar ages, race, ethnicity, and body mass index (BMI). The two asthma groups had comparable FeNO concentrations, spirometry values, total serum IgE concentrations and allergen sensitization. At the time of BEC collection, a greater proportion of the SE group were taking inhaled corticosteroids (ICS).

**Table 1.**
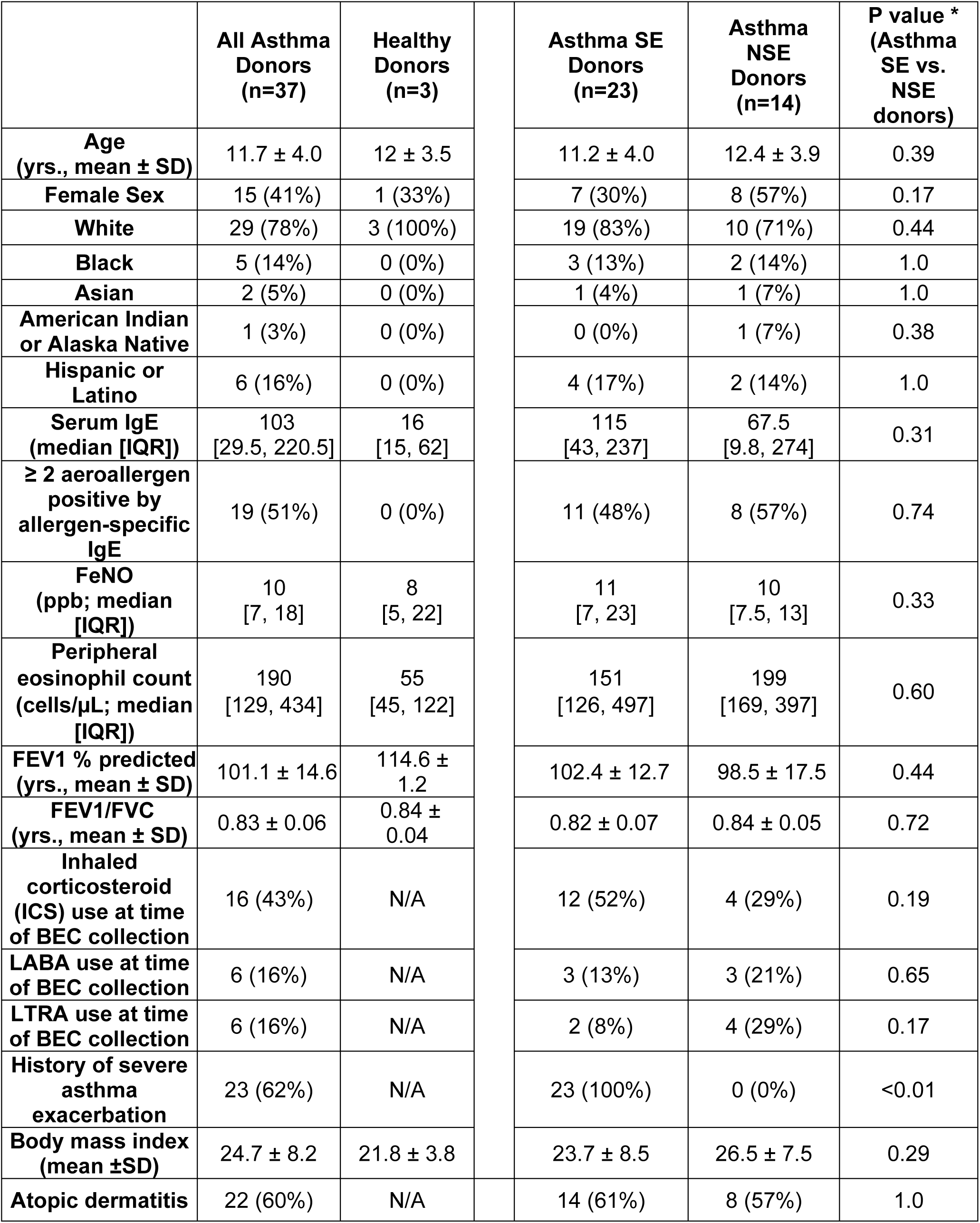
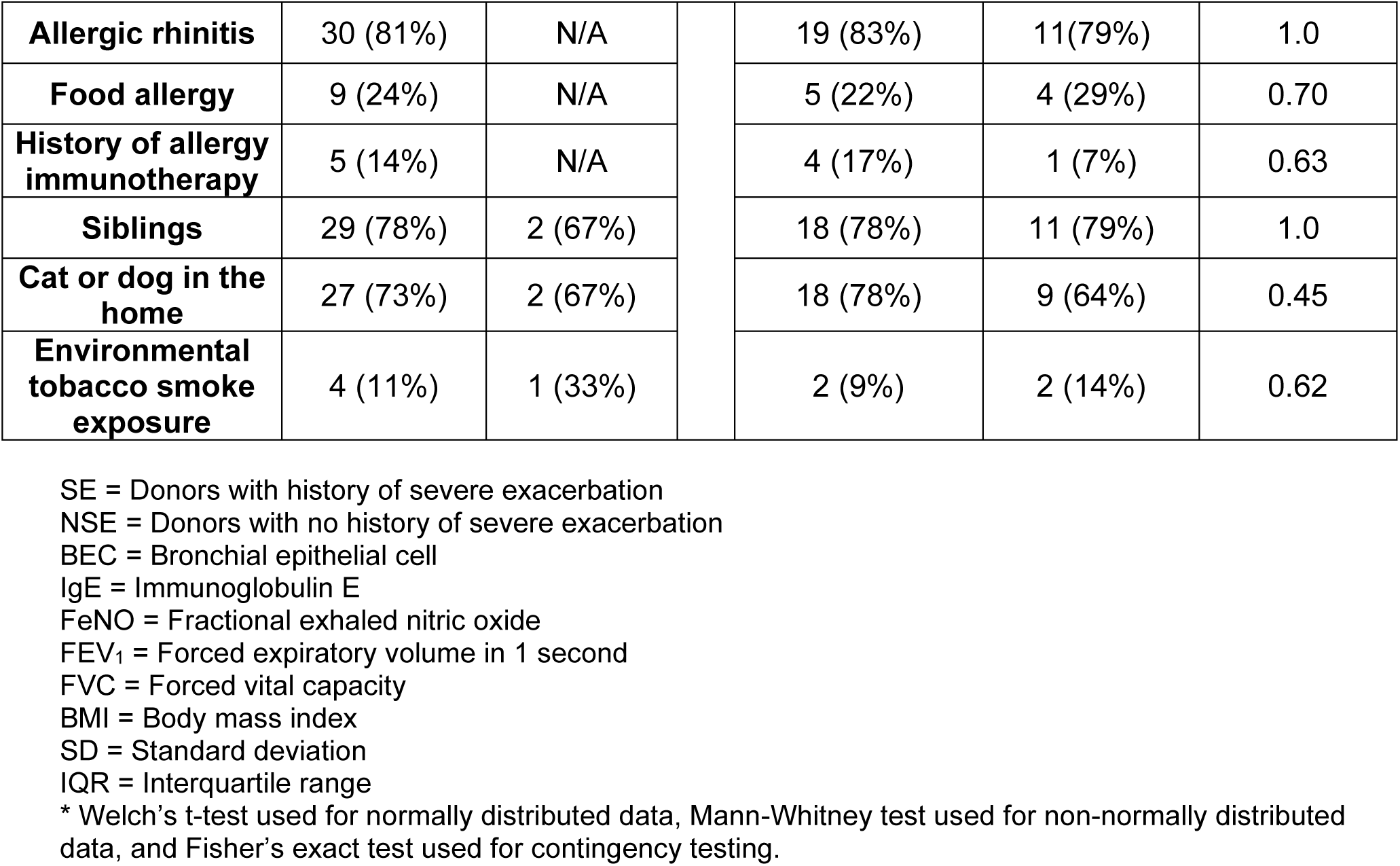
BEC donor characteristics.

|  | **All Asthma Donors (n=37)** | **Healthy Donors (n=3)** |  | **Asthma SE Donors (n=23)** | **Asthma NSE Donors (n=14)** | **P value \* (Asthma SE vs. NSE donors)** |
| --- | --- | --- | --- | --- | --- | --- |
| **Age (yrs., mean ± SD)** | 11.7 ± 4.0 | 12 ± 3.5 | | 11.2 ± 4.0 | 12.4 ± 3.9 | 0.39 |
| **Female Sex** | 15 (41%) | 1 (33%) |  | 7 (30%) | 8 (57%) | 0.17 |
| **White** | 29 (78%) | 3 (100%) |  | 19 (83%) | 10 (71%) | 0.44 |
| **Black** | 5 (14%) | 0 (0%) |  | 3 (13%) | 2 (14%) | 1.0 |
| **Asian** | 2 (5%) | 0 (0%) |  | 1 (4%) | 1 (7%) | 1.0 |
| **American Indian or Alaska Native** | 1 (3%) | 0 (0%) |  | 0 (0%) | 1 (7%) | 0.38 |
| **Hispanic or Latino** | 6 (16%) | 0 (0%) |  | 4 (17%) | 2 (14%) | 1.0 |
| **Serum IgE (median [IQR])** | 103 [29.5, 220.5] | 16 [15, 62] |  | 115 [43, 237] | 67.5 [9.8, 274] | 0.31 |
| **$\geq 2$ aeroallergen positive by allergen-specific IgE** | 19 (51%) | 0 (0%) |  | 11 (48%) | 8 (57%) | 0.74 |
| **FeNO (ppb; median [IQR])** | 10 [7, 18] | 8 [5, 22] |  | 11 [7, 23] | 10 [7.5, 13] | 0.33 |
| **Peripheral eosinophil count (cells/μL; median [IQR])** | 190 [129, 434] | 55 [45, 122] |  | 151 [126, 497] | 199 [169, 397] | 0.60 |
| **FEV1 % predicted (yrs., mean ± SD)** | 101.1 ± 14.6 | 114.6 ± 1.2 | | 102.4 ± 12.7 | 98.5 ± 17.5 | 0.44 |
| **FEV1/FVC (yrs., mean ± SD)** | 0.83 ± 0.06 | 0.84 ± 0.04 | | 0.82 ± 0.07 | 0.84 ± 0.05 | 0.72 |
| **Inhaled corticosteroid (ICS) use at time of BEC collection** | 16 (43%) | N/A |  | 12 (52%) | 4 (29%) | 0.19 |
| **LABA use at time of BEC collection** | 6 (16%) | N/A |  | 3 (13%) | 3 (21%) | 0.65 |
| **LTRA use at time of BEC collection** | 6 (16%) | N/A |  | 2 (8%) | 4 (29%) | 0.17 |
| **History of severe asthma exacerbation** | 23 (62%) | N/A |  | 23 (100%) | 0 (0%) | <0.01 |
| **Body mass index (mean ± SD)** | 24.7 ± 8.2 | 21.8 ± 3.8 | | 23.7 ± 8.5 | 26.5 ± 7.5 | 0.29 |
| **Atopic dermatitis** | 22 (60%) | N/A |  | 14 (61%) | 8 (57%) | 1.0 |
| **Allergic rhinitis** | 30 (81%) | N/A |  | 19 (83%) | 11 (79%) | 1.0 |
| **Food allergy** | 9 (24%) | N/A |  | 5 (22%) | 4 (29%) | 0.70 |
| **History of allergy immunotherapy** | 5 (14%) | N/A |  | 4 (17%) | 1 (7%) | 0.63 |
| **Siblings** | 29 (78%) | 2 (67%) |  | 18 (78%) | 11 (79%) | 1.0 |
| **Cat or dog in the home** | 27 (73%) | 2 (67%) |  | 18 (78%) | 9 (64%) | 0.45 |
| **Environmental tobacco smoke exposure** | 4 (11%) | 1 (33%) |  | 2 (9%) | 2 (14%) | 0.62 |

SE = Donors with history of severe exacerbation
NSE = Donors with no history of severe exacerbation
BEC = Bronchial epithelial cell
IgE = Immunoglobulin E
FeNO = Fractional exhaled nitric oxide
FEV<sub>1</sub> = Forced expiratory volume in 1 second
FVC = Forced vital capacity
BMI = Body mass index
SD = Standard deviation
IQR = Interquartile range
\* Welch's t-test used for normally distributed data, Mann-Whitney test used for non-normally distributed data, and Fisher's exact test used for contingency testing.

Organotypic air liquid interface (ALI) BEC cultures (Figure 1A, Supplemental Figure 1) were sampled pre-infection, infected with rhinovirus A16, and sampled at 2 days(d), 4d, 7d, and 10d post-infection. Differences in viral load levels and kinetics were compared over time among groups using both a linear model (LM) and a generalized additive mixed model (GAMM). BECs from SE donors demonstrated 5.9-fold higher viral copy number post-infection compared to NSE donors (Figure 1B, Post-infection LM: Estimate=0.77, p=2.2e-04) with higher values throughout (GAMM Shape, p=2.0e-16). This result was also significant if the models were adjusted for use of ICS or FeNO, neither of which showed significant relationship with viral load.

**Figure 1.**
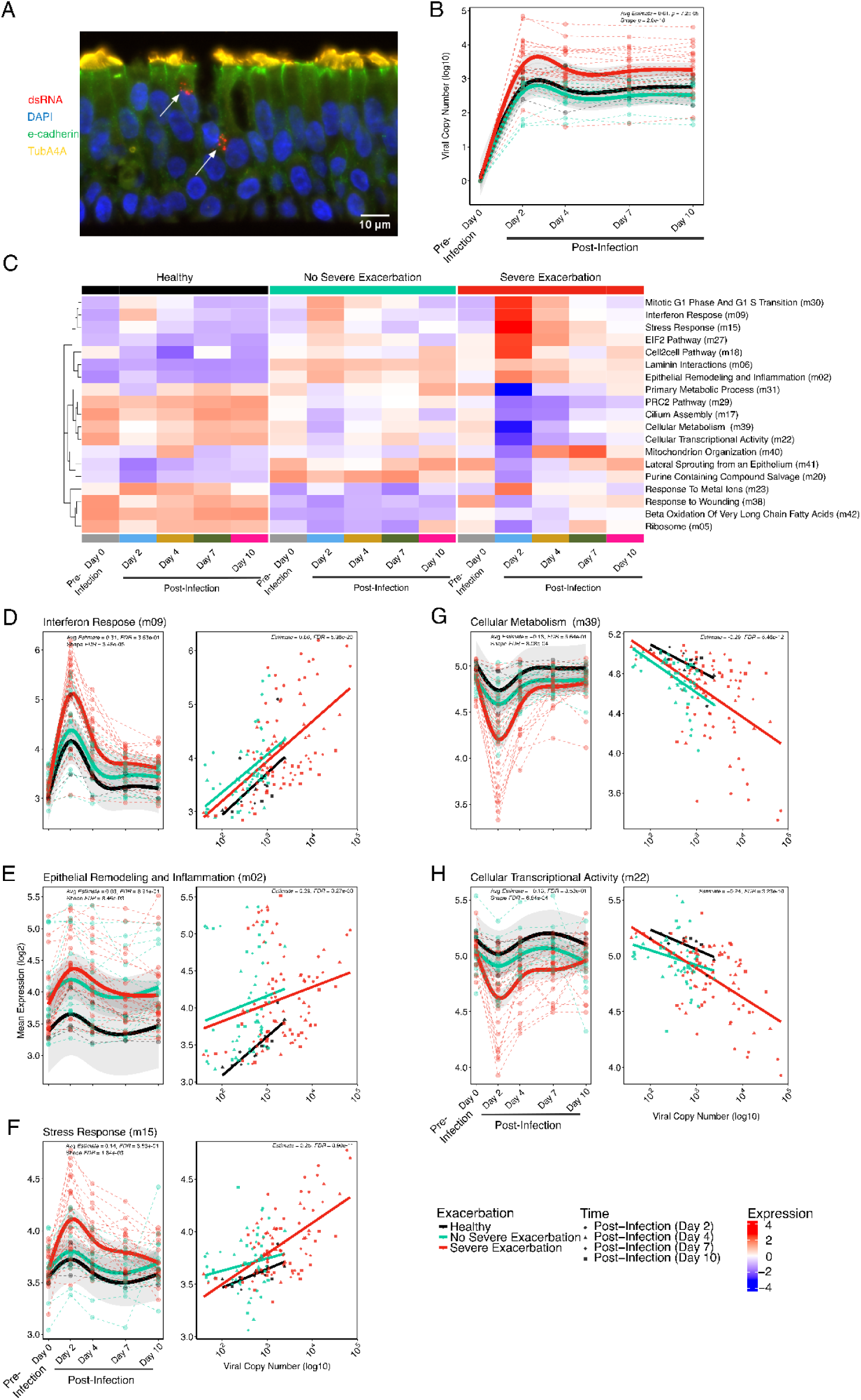
BECs from children with severe asthma exacerbations show enhanced RV replication and sustained upregulation of IFN and inflammatory and dysregulated metabolic pathways. (**A**) Immunofluorescence image of an asthma donor BEC culture 2 days after RV infection, stained with antibodies against e-cadherin (green), TubA4A (yellow), dsRNA (red). Nuclei stained with DAPI (blue). Arrows indicate replicating virus. Scale bar = 10 µm. (**B**) GAMM plot showing the non-linear changes in viral load over time differing among clinical exacerbation groups and HC. (**C**) The heat map shows the 19 significantly differentially expressed modules identified comparing the SE, NSE, and HC groups as determined by the GAMM. Module expression levels are shown as row normalized Z-scores of the mean expression for each group and timepoint (columns) with red representing higher relative expression and blue representing lower relative expression. Module row ordering is by hierarchical clustering. The five modules of highest interest and presented in the main results section are italicized. Those modules are named according to their annotation summary. The other modules are labeled by their top pathway enrichment score. The number in parentheses indicates the module number as defined by WGCNA. (**D**) Scatterplot showing the non-linear changes in expression of the “Interferon Response” module over time by donor group. Linear regression plot showing association between the “Interferon Response’’ module and viral load by donor group and time post infection (d2-10). There was a significant relationship between these two variables. (**E**) Analogous plots of the “Epithelial Remodeling and Inflammation” module demonstrating a linear relationship between module expression and viral load. (**F**) Analogous plots of the “Stress Response” module demonstrating a significant linear relationship between module expression and viral load. (**G**) Scatterplot showing the non-linear changes in expression of the “Cellular Metabolism” module over time differing by clinical exacerbation group. Regression plot showing association between the “Cellular Metabolism” module and viral load by donor group and time in the post infection timepoints (d2-10). (**H**) Analogous plots of the “Cellular Transcriptional Activity” module demonstrating a significant relationship between module expression and viral load. HC=15 samples from 3 donors, NSE=70 samples from 14 donors, SE=113 samples from 23 donors. Statistics shown indicate the SE vs NSE GAMM Shape FDR value, SE vs NSE GAMM Avg Estimate and FDR values and the all timepoint linear estimate and FDR values. Fit lines are based on a generalized additive mixed model including 95% confidence intervals.

### Children with severe exacerbations show sustained upregulation of inflammatory pathways and downregulation of metabolic pathways following RV infection

Modular analysis of RNA-seq data from BEC samples among groups over the time series (d0-d10) identified 19/42 differentially expressed modules comparing SE with NSE donors in a GAMM model (FDR<0.05; Figure 1C; Supplemental Table 1). Of these, 11/19 modules also showed significant linear associations to viral load values across the post-infection timepoints (FDR< 0.05; Supplemental Table 2). We investigated the kinetic patterns and biologic functions of these 11 modules and here highlight 5 modules of highest biological interest, with the remainder in Supplemental Table 3.

Three modules that were increased by RV infection and showed higher peak and/or greater sustained expression over time in the SE compared to NSE groups are shown in Figure 1D-F. The first module is composed of 363 genes, predominantly canonical anti-viral type I/III interferon response genes (annotated “Interferon Response”; Supplemental Figure 2). BECs from SE donors exhibited a significantly higher average expression level at d2, with a 1.69-fold increase compared to NSE donors (LM FDR=9.2e-03; Supplemental Table 4) and a 1.92-fold increase compared to HC with significantly different kinetics characterized by sustained higher levels at each post-infection timepoint among SE donors (SE vs NSE GAMM Shape: FDR=3.5e-05; Figure 1D). This module was significantly associated with viral load (Estimate=0.66; FDR=5.4e-20). It includes transcription factors involved in antiviral and inflammatory processes (*IRF7, IRF1, STAT1, STAT2*), viral signaling molecules (*CXCL10, CXCL11*), and genes with RNA-virus sensing properties (OAS family genes, *RIGI* and *IFIH1*). Interestingly it also includes smaller subclusters of Th17 inflammation genes (e.g. *IL23A* and *IL12B*) and epithelial remodeling genes (*SMAD1, CCN4*, keratin and cytoskeleton genes).

A second module increased after infection in SE is composed of 1,396 genes with densely interconnected functions representing broad cellular functions including cell development and stimulus responses. Smaller subnetworks that are significantly enriched and of particular interest included TNF-α signaling, IL-17 signaling, and extracellular matrix organization. We annotated this module “Epithelial Remodeling and Inflammation”. Expression of this module similarly peaked at 2d in all groups with non-significant 1.15-fold higher expression level at 2d in SE vs NSE (LM FDR=6.5e-01) and 1.67-fold in SE vs HC but sustained significant elevation in SE at 4d vs return to baseline levels in the NSE group (SE vs NSE: GAMM Shape: FDR=8.5e-03; Figure 1E). Module expression was significantly associated with viral load (Estimate=0.29; FDR=3.3e-03).

A third module increased in SE is annotated “Stress Response”. It includes 242 genes and is significantly enriched for unfolded protein and stress response genes (e.g. *VEGFA, NOD2, CEBPB, ATF4*), proteasome subunits, and steroid biosynthesis genes. This module was uniquely elevated in SE compared to the NSE and HC across d2-10 post-infection. SE demonstrated 1.23-fold and 1.33-fold higher expression at 2d compared to NSE (LM FDR=1.3e-02) and HC, respectively, and a sustained increase over time in SE (SE vs NSE: GAMM Shape: FDR=1.8e-3; Figure 1F). Expression of this module was also significantly associated with viral load (Estimate=0.25; FDR=8.9e-11).

Two modules that were decreased by RV infection and showed a lower nadir and lower sustained expression over time in SE compared to NSE and HC are shown (Figure 1G, H). The first we annotated “Cellular Metabolism”, includes 106 genes, and is significantly enriched for genes related to lipid oxidation, branched-chain amino acid degradation (e.g. *HADHB*, *MCCC1*, *ACADL*), and oxidative phosphorylation. Interestingly, this module also contains several genes related to beta-adrenergic-receptor signaling and muscarinic-acetylcholine receptor signaling (*SNAP29*, *PRKX*, *ALDH6A1*). The nadir of expression of this module occured at d2 showing 1.30-fold and 1.52-fold lower expression in SE vs NSE (LM FDR=1.3e-02) and HC, respectively. This module remained lower in SE through the later timepoints, (SE vs NSE: GAMM Shape: FDR=8.1e-04). Expression of this module was inversely associated with viral load (Estimate=-0.29; FDR=5.5e-12).

The second such module includes 204 genes, the largest subset relevant to cellular transcriptional activity, a smaller subset related to amino acid metabolism; we annotated this “Cellular Transcriptional Activity”. This module also nadired at d2 showing 1.21-fold and 1.32-fold lower expression in SE vs NSE (LM FDR=1.28e-02) and HC, respectively, with expression remaining lower in SE asthma through d10 (SE vs NSE: GAMM Shape: FDR=6.8e-04). Expression of this module was also inversely associated with viral load (Estimate=-0.28; FDR=3.2e-11).

A linear model subset to the post-infection timepoints, d2-10, was used to determine the extent to which observed viral load differences statistically mediated the observed differences in module expression between the SE and NSE groups for the 5 modules of interest. We observed that for the Interferon Response, Stress Response, Cellular Metabolism, and Cellular Transcriptional Activity modules viral load mediated most of the difference in module expression observed between the two groups (Supplemental Figure 3; Supplemental Table 5).

### Differences in pre-infection IFN tone relate to the observed greater viral replication in children with severe exacerbation

We investigated pre-infection differences in IFN tone between BEC cultures from SE and NSE that might explain the observed differences in viral infection kinetics. While there was no pre-infection difference in type I/III IFN gene expression between SE and NSE at a stringent FDR<0.05, gene set enrichment analysis (GSEA) demonstrated lower aggregate expression of the Hallmark IFN alpha response pathway in SE compared to NSE (Figure 2A). We measured protein concentrations of interferon-beta (IFNβ), interferon-lambda-2 (IFN-λ2/IL-28A), interferon-lambda-3 (IFN-λ3/IL-28B), and the interferon-stimulated chemokine CXCL10 in supernatant from BECs pre-infection and 2-days following RV infection. All 3 IFNs were below the assay detection limit in BECs pre-infection, IFN-β remained below the assay detection limit in most samples following RV infection while IFN-λ2 and IFN-λ3 were consistently detected following RV infection. In contrast, CXCL10 was readily detected with broad dynamic range in pre- and post-infection samples and was therefore used as a surrogate protein marker of IFN tone. We observed that pre-infection secreted CXCL10 protein concentrations were 3.32-fold lower in SE compared to NSE (p=6.9e-04) and 2.88-fold lower compared to HC (p=1.0e-01; Figure 2B and Supplemental Table 6). Furthermore, pre-infection BEC CXCL10 protein concentration was significantly associated with the subsequent post-infection viral load at d2 in an inverse log-linear relationship (d2; Pearson R=-0.38, p=1.6e-02; Figure 2C; Supplemental Figure 4) and was also inversely associated with secreted IFN-λ2, IFN-λ3 and CXCL10 protein at d2 following RV infection (Supplemental Figure 5).

**Figure 2.**
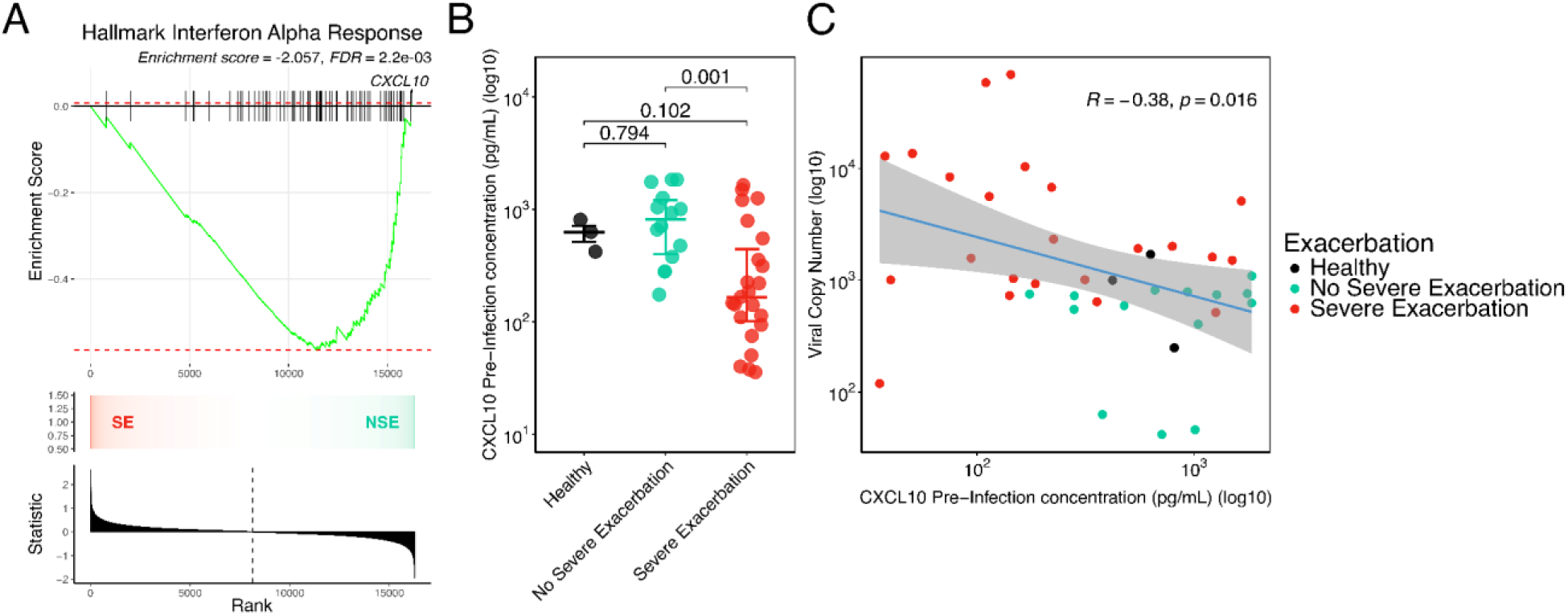
Pre-infection IFN tone drives viral replication in children with severe exacerbation. (**A**) GSEA analysis shows a significantly negative enrichment score of the MSigDB Hallmark IFN alpha response geneset in the SE group compared to the NSE group in the pre-infection d0 samples (GSEA Enrichment score=-2.057, FDR=2.2e-03). The plot shows the running enrichment score for the gene set from decreasing values of the differential expression rank list. The peak represents the ES for the gene set. The bars show the genes in the geneset ranked by effect size relative to all genes which are shown in the ranked list metric plot. The CXCL10 gene has been indicated. The color gradient indicates which comparison group the genes appear in the ranked list of genes.The bar plot at the bottom represents the effect size of all gene ranking. (**B**) Dot plot showing log transformed pre-infection CXCL10 concentration values by exacerbation status, boxplots indicate median and interquartile ranges for each group. (**C**) Scatter plot showing a significant inverse relationship between log transformed pre-infection CXCL10 values and log transformed viral load at d2 (Pearson R = - 0.38, p=1.6e-02). Fit line is based on a linear model including 95% confidence intervals. HC n=3 donors, NSE n=14 donors, SE n=23 donors.

### Prophylactic IFN-β treatment reduces RV replication and ameliorates post-infection epithelial IFN, inflammatory, remodeling, and metabolism dysregulation

To determine whether low IFN tone and/or low CXCL10 levels pre-infection are causal factors promoting RV replication and the differential post-infection inflammatory responses observed dysregulated in SE donors, we treated BEC cultures with IFN-β or CXCL10 starting 2 days prior to RV infection and continued for 2 days after RV infection. IFN-β treatment led to a significant decrease in viral load at 2d compared to the untreated samples in both asthma groups with 4.22-fold decrease in the SE group (p=4.81e-04) compared to a 3.39-fold decrease in the NSE group (p=1.7e-04) (Figure 3A). Treatment of BEC cultures with CXCL10 did not impact viral load (data not shown). IFN-β treatment also resulted in a 1.53-fold relative decrease in expression of the Interferon Response module at 2d (Estimate=-0.61, FDR=1.2e-03; Figure 3B; Supplemental Table 7). Regression analysis demonstrated that the 2d viral load had direct association with the expression of this module (β=0.51, p=3.1e-86; Supplemental Table 8), indicating IFN-β treatment resulted in lower viral load, which significantly and completely mediated the relatively diminished Interferon Response module expression (Supplemental Table 9).

**Figure 3.**
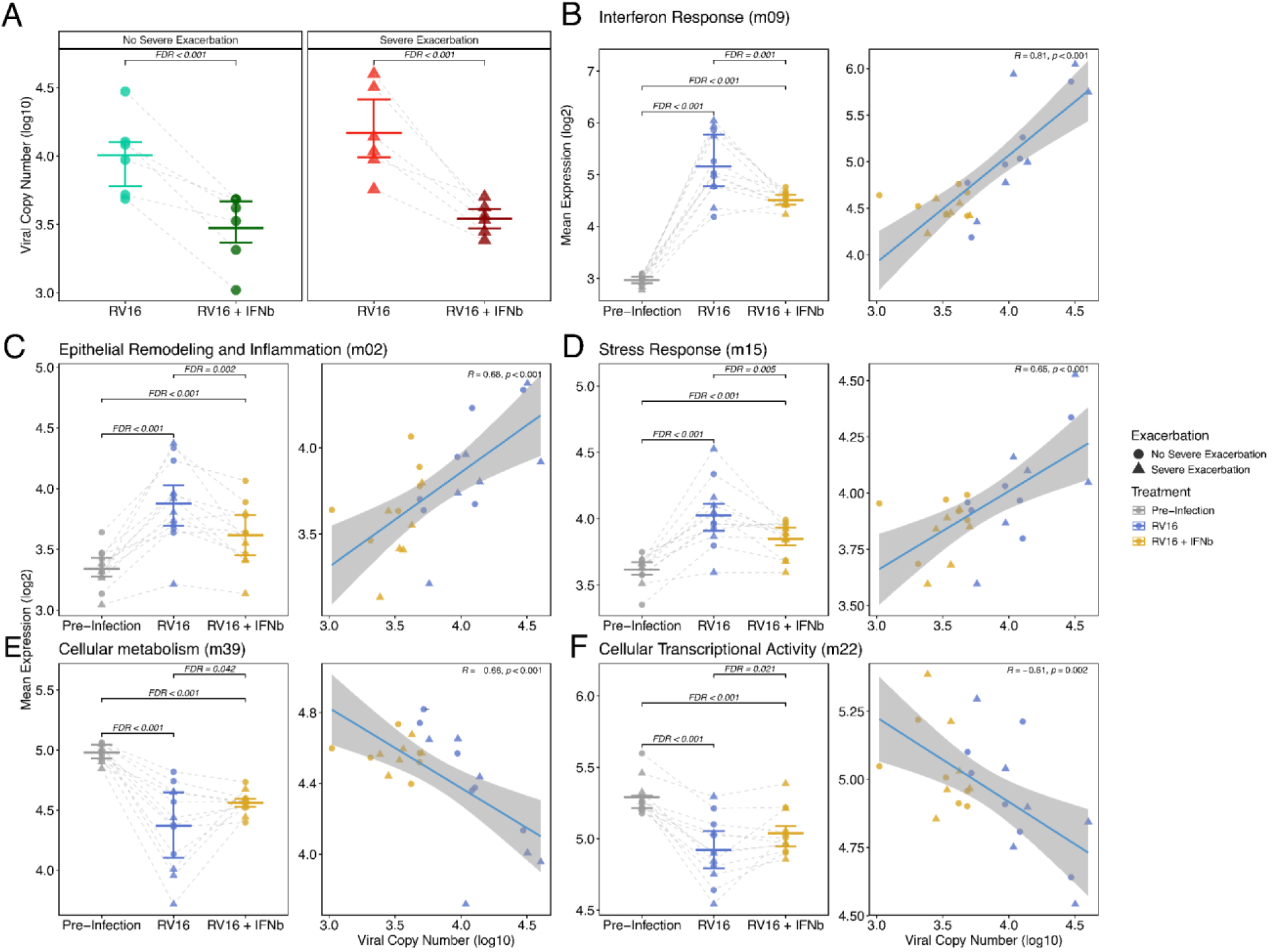
Prophylactic IFN-β treatment reduces RV replication and ameliorates epithelial IFN, inflammatory and remodeling pathways. (**A**) Dot plot showing log transformed viral copy number values by exacerbation status between samples at Day 2 treated with IFN-β. The IFN-β treated group showed a significant decrease in viral load compared to the Untreated samples in both exacerbation groups with a 3.39-fold decrease in the NSE group (n=6 donors) and a 4.22-fold decrease in the SE group (n=6 donors), boxplots indicate mean and interquartile ranges for each group. (**B**) Dot plot showing expression changes in Interferon response module by IFN-β treatment. Regression plot showing association between the “Interferon response” module and viral load by IFN-β treatment. (**C**) Dot plot showing expression changes in Epithelial Remodelling and Inflammation Response module by IFN-β treatment. Regression plot showing association between the “Epithelial Remodelling and Inflammation Response” module and viral load by IFN-β treatment. (**D**) Dot plot showing expression changes in Stress response module by IFN-β treatment. Regression plot showing association between the “Stress response” module and viral load by IFN-β treatment. (**E**) Dot plot showing expression changes in Cellular metabolism module by IFN-β treatment. Regression plot showing association between the “Cellular metabolism” module and viral load by IFN-β treatment. (**F**) Dot plot showing expression changes in Cellular transcriptional activity module by IFN-β treatment. Regression plot showing association between the “Cellular transcriptional activity” module and viral load by IFN-β treatment. Boxplots indicate median and interquartile ranges for each group and the Fit line is based on a linear model including 95% confidence intervals.

The Epithelial Remodeling and Inflammation module showed a 1.20-fold relative decrease in expression following IFN-β treatment (Estimate=-0.27, FDR=2.2e-03), with significant positive association with viral load (β=0.12, FDR=8.2e-19; Figure 3C), which also showed significant mediation. The Stress Response module exhibited a 1.14-fold relative decrease in expression with IFN-β treatment (Estimate=-0.19, FDR=4.6e-03) and significant association with viral load (β=0.10, FDR=6.6e-19; Figure 3D) and significant mediation. In contrast, CXCL10 treatment did not impact expression of the Interferon Response, Epithelial Remodeling and Inflammation, or Stress Response modules (Supplemental Figure 6).

IFN-β treatment also led to a relative restoration in expression levels of modules that had been significantly decreased in SE donors following infection. The Cellular Metabolism module displayed a 1.12-fold relative increase in expression with IFN-beta treatment (Estimate=0.16, FDR=4.2e-02) and an inverse relationship with viral load (β=-0.13, FDR=4.6e-25, Figure 3E) and significant mediation. Similarly, the Cellular Transcriptional Activity module showed a 1.09-fold relative increase in expression (Estimate=0.13, FDR=2.1e-02), an inverse association with viral load (β=-0.09, FDR=3.4e-24, Figure 3F), and significant mediation. CXCL10 treatment did not impact expression of either module.

### scRNA-sequencing confirms and demonstrates the cellular sources of dysregulated responses to RV in children with severe exacerbation

To investigate epithelial cell type specific responses in BECs pre- and post-RV infection we generated scRNA-seq data from a subset of donors in an independent experiment: SE (n = 7), NSE (n = 4) and healthy (n = 3). These yielded 316,712 individual cells after filtering and quality control. Unsupervised, graph based clustering identified 12 coarse cell clusters, which we annotated using cell-type specific markers for airway epithelial cells (12–14) (Figure 4A). This analysis revealed expected major epithelial cell populations as well as a unique population with the highest pre- and post-RV expression of a diverse array of inflammatory genes such as *ISG20, IFIH1, IFIT1, CXCL1, CXCL3, CXCL5, CXCL8, CXCL10* (Figure 4B; Supplemental Table 10), alongside expression of markers typical of suprabasal, secretory, and goblet cells, consistent with a secretory lineage. We interpret these to be secretory lineage cells involved in coordinating and/or propagating an innate immune response consistent with some past observations (15) and label them secretory immune response cells (SIRs). We identified these 12 cell populations in each sample/donor without any sample bias (Figure 4C; Supplemental Table 11); there were no significant differences in cell composition among donor groups (SE, NSE and HC) or by pre- and post-infection status. Custom viral probes were generated for the experiment and confirmed infection but showed relatively low hybridization preventing statistically robust assessment of viral quantity at the single cell level (see Methods).

**Figure 4.**
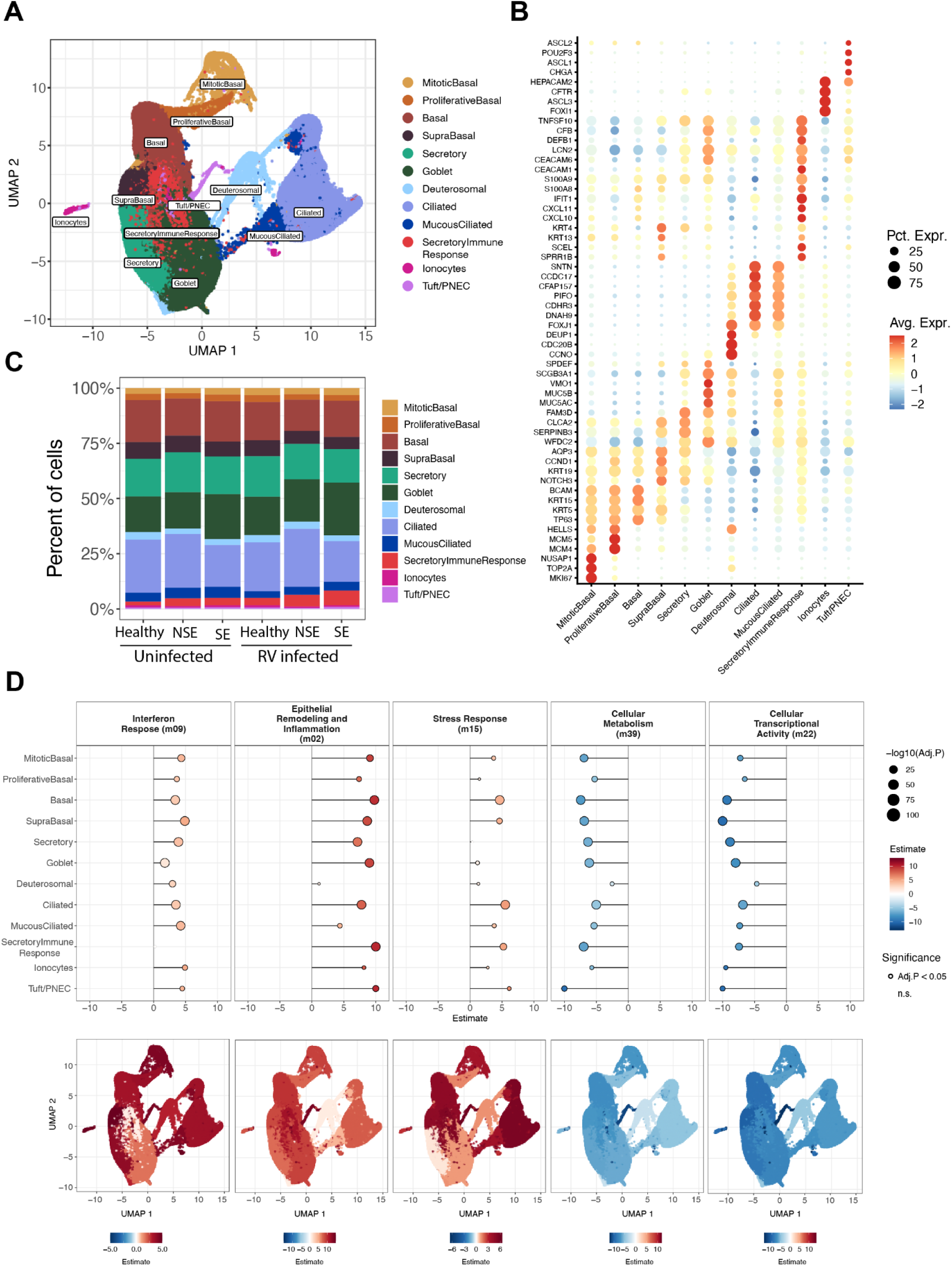
scRNA-seq maps epithelial cell diversity and reveals cell type–specific activation of bulk-derived transcriptional modules following RV infection. (**A**) UMAP representation of the 316,712 cells recovered from samples collected from children with asthma (SE = 7 donors, NSE = 4 donors) and healthy controls (n = 3 donors). Mitotic Basal: 8,281 cells, Proliferative Basal: 8,405 cells, Basal: 53,663 cells, Supra Basal: 20,403 cells, Secretory: 52,727 cells, Goblet: 63,542 cells, Inflammatory: 14,361 cells, Deuterosomal: 8,836 cells, Mucous-Ciliated: 13,284, Ciliated: 67,654 cells, Ionocytes: 2,407 cells, Tuft/PNECs: 2,251 cells. (**B**) The dot plot shows the expression of marker genes distinguishing distinct cell populations from the EmptyDrops based processing pipeline. The color intensity indicates the magnitude of marker gene expression, and the size of the circle denotes percent of cells expressing the marker gene. (**C**) Bar plot showing the mean proportions of identified epithelial cell populations by scRNA-seq by condition (uninfected and 2 days post-infection samples) and donor groups (SE = 7 donors, NSE = 4 donors and healthy controls = 3 donors). (**D**) The dot plot showing the CLM estimates for select modules in RV infected samples across all 12 cell types. The corresponding bottom plots show the relative CLM estimates on a UMAP space.

We investigated cell-level expression differences among donor groups in post-infection samples for the same modules derived from the bulk RNA-sequencing analysis. We used ordinal regression to identify those modules with either a significant ascending (SE > NSE > HC) or descending expression trend (SE < NSE < HC) among donor groups (Supplemental Table 12). In the post-infection samples, the Interferon Response module was highest in SE asthma showing an ascending trend in all but one cell type (Figure 4D, Supplemental Figure 7), consistent with the bulk transcriptomics data. The magnitude of this ascending difference among donor groups varied among cell types. Curiously this module showed no expression difference among groups in the SIRs (FDR>0.05), which showed the highest overall expression of this module. Goblet cells showed relatively modest differences among groups (Estimate=1.7, FDR= 4.7e-120). All other cell types showed fairly consistent large differences (Estimates=3.0-4.9, FDR<10e-10). Expression of this module was markedly increased in post-infection compared to pre-infection samples in all cell types and donor groups.

Expression of the Epithelial Remodeling and Inflammation module also showed an ascending trend in post-infection samples in all cell types. Magnitude varied markedly by cell type (Estimates=1.1-13.9). This module expression difference by group was particularly pronounced in the tuft/pulmonary neuroendocrine cells (PNECs) (Estimate=13.9, FDR<10e-10) and SIRs (Estimate=11.6, FDR<10e-10). It was low in deuterosomal cells (Estimate=1.1, FDR=1.6e-02). The Stress response module showed an ascending trend in post-infection samples (Estimates=1.2-6.1). This was particularly pronounced in tuft/PNEC cells (Estimate=6.1, FDR=4.9e-10), ciliated cells (Estimate=5.5, FDR<10e-10), and SIRs (Estimate=5.2, FDR<10e-10) but low in goblet cells (Estimate=1.2, FDR=1.3e-09) and not significant in secretory cells.

The Cellular Metabolism and Cellular Transcriptional Activity modules showed descending expression trends (SE < NSE < HC) in post-infection samples consistent with the bulk transcriptomics data. Both modules showed similar trends among cell types, with differences particularly pronounced in tuft/PNEC and basal cells (Estimates=-7.4 to -13.9, FDR<10e-10), and least in deuterosomal cells (Estimates=-2.5 to -4.7, FDR<10e-10).

Further analysis of gene level expression differences among the three donor groups was performed to identify additional cell specific signals that might not have been detected in the module level bulk transcriptomics data. In the post-infection samples, 5,160 unique genes showed an ascending trend among donor groups in one or more cell types while 5,659 unique genes had a descending trend (Figure 5A; Supplemental Table 13). The SIR cell cluster had the greatest transcriptional difference among donor groups, (4,779 genes ascending; 488 descending). Genes with an ascending trend in the SIR cell cluster were a broad array of inflammatory molecules such as *IL1B*, *CXCL5*, *LCN2*, *FN1*, *LAMA4*, *CLIC5*, *GATA3*, and *ALOX5* (Figure 5B). The gene list was enriched for pathways related to lysosomes, focal adhesion, neutrophil extracellular trap formation, and antimicrobial response pathways.

**Figure 5.**
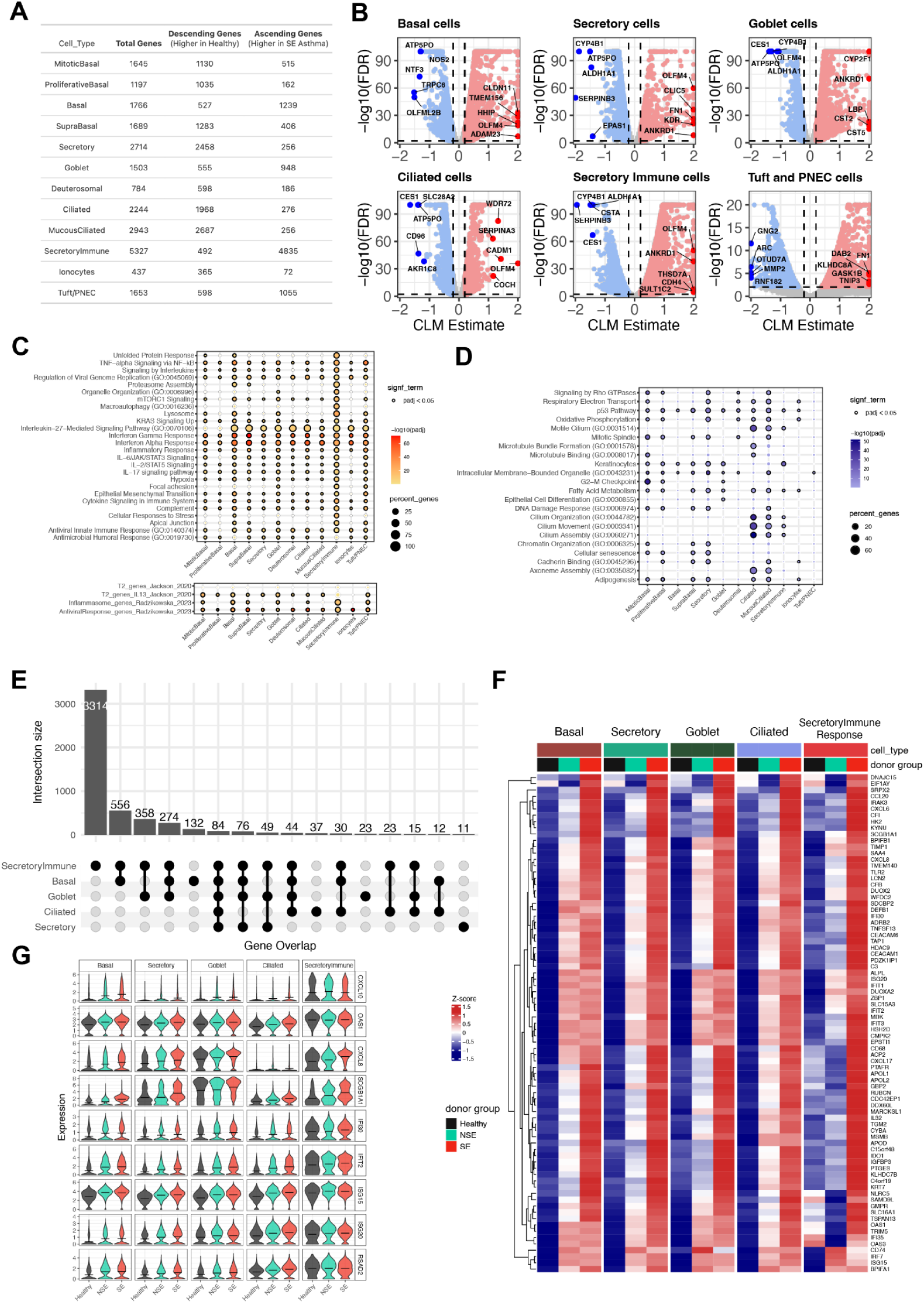
Global gene expression analysis identifies severity-associated transcriptional programs across epithelial cell types following RV infection. (**A**) The count of genes with ascending (SE > NSE > HC) and descending (SE < NSE < HC) expression trend across donor groups (SE = 7 donors, NSE = 4 donors, healthy controls = 3 donors) in each cell type. (**B**) The volcano plots show the genes with an ascending and descending trend across donor groups in basal cells, secretory cells, goblet cells, ciliated cells, and secretory immune response cells. Significant (FDR < 0.01) genes with an ascending trend (estimate > 0.2) and descending trend (estimate < -0.2) that are expressed in at least 10% of cells are denoted by red and blue color points, respectively. Select top ranked genes by estimate are highlighted. (**C**) Dot plot showing enriched GO terms and pathways on significant genes with an ascending trend. The size of each dot denotes the percentage of genes, and the intensity of color denotes statistical significance as –log10(adj.*P*). (**D**) Dot plot showing enriched GO terms and pathways on significant genes with descending trend. The size of each dot denotes the percentage of genes, and the intensity of color denotes statistical significance as –log10(adj.*P*). (**E**) Upset plot showing the count of overlapping genes with an ascending trend across five cell types, basal, secretory, goblet, ciliated and secretory immune response cells. (**F**) Heatmap showing relative expression of 84 core signature genes (shared across all five cell types) with an ascending expression trend at 5 cell types. Gene expression levels are shown as row normalized Z-scores of the mean expression for each group with red representing higher relative expression and blue representing lower relative expression. (**G**) Violin plot showing the distribution of gene expression for select genes with an ascending trend across donor groups by individual cell types. The black horizontal line in each violin denotes a mean expression.

To globally characterize the differences among donor groups by cell type in this gene level analysis, we performed pathway enrichment of the ascending and descending genes. Multiple immune, stress, and remodeling pathways such as interferon responses, IL-6/JAK/STAT3 signaling, IL-17 signaling, TNF-alpha, and hypoxia, were enriched among those ascending genes highest in SE, particularly in SIRs, basal cells, tuft/PNECs, and goblet cells (Figure 5C; Supplemental Table 14). The asthma associated pan-epithelial remodeling of IL-13 gene signature described by Jackson et al. was particularly enriched among ascending genes in goblet and basal cell populations (16). The asthma associated inflammasome gene signature described by Radzikowska et al. was particularly enriched among ascending genes in SIRs and basal cell clusters (17). Among descending genes, lowest in SE, cilium assembly and related pathways were particularly enriched in ciliated and mucous ciliated cells, while transcriptional regulation and energy metabolism pathways were particularly enriched in secretory cells (Figure 5D; Supplemental Table 14). We also identified a “core gene signature” with consistent trend (same directionality and significant) and shared across five major cell populations, basal, secretory, goblet, ciliated and SIR cells. This included 84 genes showing an ascending trend across donor groups (SE > NSE > HC) inclusive of viral response genes *RSAD2*, *ISG15*, *IFI30*, *IFIT2*, *OAS1*, *STAT1* and *ZBP1,* chemokines and inflammation markers *CXCL6*, *CXCL8*, *CXCL10*, *IL32* and complement related genes *C3*, *CFI*, *APOL1* and *APOL2* (Figure 5E-G). It included 48 genes with a descending trend across donor groups (SE < NSE < HC) including epithelial barrier function genes (*UPK1B*, *EPPK1*, *EVPL*) and antioxidant response genes (*GCLC, SELENOP*) (Supplemental Figure 8).

## Discussion

We have investigated kinetics of RV replication and host responses in organotypic bronchial epithelial cultures from well-characterized asthmatic and healthy children using bulk and single-cell RNA sequencing and targeted protein assessment. We found higher viral loads in cultures from asthmatic children with a history of severe exacerbations compared to those without or healthy controls. Transcriptomic modular analysis showed heightened and prolonged expression in SE donors of interferon response genes, inflammatory and remodeling pathways, and unfolded protein stress responses, all of which positively correlated with viral load. The Interferon Response module included not only canonical ISGs but also genes involved in Th17 inflammation and epithelial remodeling, which together with the Epithelial Remodeling and Inflammation module, illustrate how impaired antiviral control may converge on broader pro-inflammatory and tissue-altering pathways in exacerbation-prone epithelium. In contrast, pathways related to cellular metabolism and transcriptional activity were downregulated in SE and inversely correlated with viral load. This cascade was evident in both bulk and single-cell analyses, with viral load statistically mediating the magnitude of these transcriptional responses and the differences observed in SE donors. Single-cell RNA-seq further revealed how several epithelial subtypes contributed differentially to these amplified responses including basal, secretory immune response, and tuft/PNECs showing particularly exaggerated post-infection activity.

Pre-infection samples from SE donors showed modestly lower type I interferon gene expression and reduced CXCL10 protein levels, which was inversely associated with post-infection viral load in a log-linear manner. This indicates that subtle deficits in baseline epithelial IFN tone confer susceptibility to greater RV replication, and downstream dysregulation. To test causality, we treated cultures with IFN-β before and during infection, which reduced viral replication and attenuated downstream transcriptional responses including ISGs themselves. These findings prove how low baseline interferon tone can permit enhanced viral replication and resultant exaggerated inflammatory responses in airway epithelium from severe exacerbation-prone children with asthma.

These findings provide mechanistic insight into how bronchial epithelial dysfunction can predispose children with asthma to severe exacerbations following RV infection. While prior studies have reported conflicting evidence regarding impaired versus exaggerated IFN responses in asthmatic epithelium (3–6, 18), our integrated approach helps reconcile these discrepancies. We show that children with a history of severe exacerbation exhibit reduced basal interferon tone, which permits greater viral replication and, in turn, provokes amplified secondary interferon, inflammatory, stress, and remodeling responses. These findings refine our understanding of viral-triggered asthma exacerbations in children by demonstrating that deficient and exaggerated interferon responses can coexist in the same epithelium, but at distinct stages. While earlier studies emphasized impaired IFN responses in asthma (3, 4), more recent work, including Bhakta et al. and Altman et al., has linked excessive ISG expression to more severe disease and reduced lung function (7, 8). Our data reconcile these views, showing that children prone to exacerbation exhibit low baseline IFN tone that permits enhanced viral replication, which in turn drives an amplified secondary IFN, inflammatory, and remodeling response. This temporal duality helps explain prior conflicting findings and highlights the importance of assessing epithelial responses dynamically. Our results underscore and help explain recent clinical findings by Gaberino et al. which showed that lower pre-infection airway ISG expression predicted both higher risk of exacerbation and greater IFN upregulation during virus-triggered exacerbation events (10).

The broader upregulation of the Epithelial Remodeling and Inflammation and Stress Response modules in SE donors is notable. These modules included networks related to IL-17, TNF, and inflammasome signaling, all of which have been implicated in non-T2 or "neutrophilic" asthma endotypes (19–23). Although these pathways have often been studied in the context of pollutant exposure or bacterial infection (24–27), our findings indicate that RV alone can induce this pro-inflammatory, non-eosinophilic, airway remodeling milieu in SE epithelium, a finding recapitulated in the single cell data. While biologics targeting IL-17 and TNF have failed to show benefit in unselected asthma populations (28, 29); perhaps only a subset of patients – e.g. those prone to severe RV-triggered exacerbations – may derive benefit from these treatment strategies. We also show that the unfolded protein response (UPR) and endoplasmic reticulum (ER) stress pathways are disproportionately activated in asthmatic children with SE. This aligns with prior observations in human asthma and animal models implicating maladaptive UPR and ER stress in epithelial dysfunction, goblet cell metaplasia, and airway remodeling (30–33). As UPR signaling has been shown to promote mucus production and viral replication (34, 35), therapeutic targeting of ER stress pathways could represent an additional strategy to mitigate post-infection epithelial injury in asthma. Strikingly, our single-cell data show that these transcriptional programs are active across multiple epithelial cell types but are notably pronounced in several non-ciliated populations – such as secretory immune response, tuft/PNEC, and basal cells – highlighting the emerging recognition of the importance of these cells in airway biology from a rapidly growing body of airway single-cell research (15, 36). Their exaggerated activation in SE donors suggests that epithelial cell-intrinsic programs – not merely infiltrating immune cells – are central to the pathophysiology driving viral-triggered asthma exacerbations. This pattern may reflect underlying genetic and/or epigenetic influences and aligns with prior studies showing that asthma-associated genetic risk variants and epigenetic modifications are enriched in genes expressed by airway epithelial cells (37–39).

Finally, our observation that IFN-β pre-treatment suppressed viral load and normalized downstream transcriptional responses strongly suggests a causal role for low basal IFN tone in predisposing to RV-induced exacerbations and epithelial dysfunction. While prior clinical trials of inhaled IFN-β in unselected asthma populations and initiated after symptom onset showed limited efficacy (40), our data argue for a prophylactic strategy targeted toward individuals with documented low IFN tone, administered early in the course of infection or during high-risk seasons. Such a precision-based approach, guided by baseline epithelial immune profiling, could improve clinical outcomes with an existent and safe potential therapeutic while reducing unnecessary treatment exposure.

Together, our results offer a comprehensive model in which deficient pre-infection IFN signaling in asthmatic bronchial epithelium permits RV replication, triggering exaggerated inflammatory, remodeling, and stress responses that contribute to the risk and severity of exacerbation. Our model integrates bulk and single-cell transcriptomic data and protein assessment, aligns with prior mechanistic and clinical observations, and identifies novel epithelial cell populations and pathways as potential therapeutic targets. In doing so, it refines current paradigms of RV-triggered asthma exacerbations by linking impaired basal antiviral tone with subsequent epithelial remodeling and pro-inflammatory cascades, a connection only incompletely captured in prior studies. Future studies exploring the genetic and epigenetic underpinnings of low IFN tone and evaluating prophylactic immunomodulatory strategies in at-risk children with asthma are warranted.

There are several limitations to our study. Our post-infection sampling began at 2 days and therefore lacked earlier timepoints that might provide increased temporal resolution of the observed inflammatory cascade; this was informed by prior work showing peak immune responses and viral replication 48–96 hours post-infection in organotypic ALI cultures (8) and consistent with in vivo observations (10). We used viral RNA copy number as a practical, cost effective, and reproducible measure of viral load instead of trying to quantify infective virions; this was due to both technical and cost limitations of conducting TCID50 or plaque assays from 248 unique ALI culture samples wherein accurate reproducible sampling from the apical surface would have been impaired by mucus layers of organotypic ALI cultures. Our small sample size of HC donors is a limitation, as larger numbers of donors would be needed to fully characterize the heterogeneity of transcriptomic responses to RV infection in healthy children. In this study, our design and statistical power were focused on the primary comparison between SE and NSE asthma groups, with HCs included as a comparative reference point rather than for hypothesis testing. Assessing donor serum antibody levels against type I/III IFNs as well as RV-A16 would provide valuable complementary information and would be an interesting future research direction that could help further elucidate mechanisms underlying differences between SE, NSE, and control groups. Unfortunately, these measurements were not performed in the current study, and we lack residual serum samples from subjects to measure serum antibody levels in this cohort. Although a highly unique strength of our study is the ability to isolate bronchial epithelial responses in a robust sample size of carefully characterized children with and without asthma, the responses we observed reflect only epithelial responses lacking direct interaction with other immune cells that would be relevant in a clinical context. These findings suggest that disease-related epithelial programs are at least partly preserved in culture, underscoring both the utility and limitations of ALI models and the need to integrate in vivo studies to fully capture epithelial response heterogeneity in asthma. RNA analysis of primary cells obtained at the time of brushing could provide valuable complementary information by linking the *in vivo* epithelial state to the transcriptional and epigenetic programs maintained in culture. Although not performed here, such paired analyses represent an important future direction to determine which signatures observed in ALI cultures most closely mirror transcriptional memory from the original epithelial progenitors.

In conclusion, bronchial epithelial cultures from asthmatic children with a history of severe exacerbations exhibited greater RV replication and exaggerated interferon, inflammatory, remodeling, and stress responses, alongside reduced metabolic and transcriptional activity, compared to NSE and healthy donors. These differences were strongly associated with lower pre-infection IFN-stimulated gene expression and CXCL10 protein levels. Prophylactic IFN-β treatment reduced viral replication and normalized downstream responses, supporting low baseline IFN tone as a causal factor in viral susceptibility and epithelial dysregulation. These findings highlight resting epithelial IFN tone as a key determinant of host response heterogeneity and suggest that early, targeted immunomodulatory interventions may help prevent severe virus-induced exacerbations in susceptible children (40, 41).

## Methods

### Sex as a biological variable

Both male and female participants were included in this study. Sex was recorded at the time of enrollment and included as a covariate in all relevant statistical analyses. Where appropriate, analyses were stratified by sex to examine potential sex-specific effects. No sex-based exclusion criteria were applied, and all procedures were performed uniformly across sexes.

### Study population and air-liquid interface (ALI) cultures of bronchial epithelial cells

Bronchial epithelial cells (BECs) from children with asthma (n=37) and healthy children (n=3) ages 6-18 years of age were obtained from subjects while under general anesthesia for an elective surgery using 4-mm Harrell unsheathed bronchoscope cytology brushes (CONMED®) inserted through an endotracheal tube as we have previously described (8, 42). Cells were then seeded onto T-25 cell culture flasks (Corning®) precoated with type I collagen and proliferated under submerged culture conditions using PneumaCult™EX-Plus medium (Stemcell™) (8). BECs (passage 2 or 3) were proliferated on transwells under submerged culture conditions using PneumaCult™EX-Plus medium (Stemcell™) for 4-7 days then differentiated to an organotypic pseudostratified ciliated state (Supplemental Figure 1) for 21 days at ALI using PneumaCult™ALI medium (Stemcell™) as previously described (8, 43–45). We have previously demonstrated stability of gene expression in primary BEC ALI cultures between passages 1 and 3 (46).

### Clinical characterization of BEC donors

BEC donors were carefully clinically characterized. Blood was drawn for total serum IgE, eosinophil count and allergen-specific IgE to common aeroallergens (cat, dog, D. farinae, D. pteronyssinus, alternaria tenuis, aspergillus fumigatus, and timothy grass). Subjects completed a clinical phenotyping visit that included: 1) a detailed review of medical and social history (e.g. asthma exacerbation history, medication use, environmental tobacco smoke exposure, cat and dog exposure, siblings in home), 2) review of atopy history (past skin prick test or allergen-specific IgE testing, physician-treated atopic dermatitis, allergic rhinitis, and food allergy, treatment, and history of allergy immunotherapy), 3) measurement of total IgE and an aeroallergen specific IgE panel, 4) performance of spirometry and 5) measurement of the fraction of exhaled nitric oxide (FENO). Spirometry was performed in accordance with American Thoracic Society/European Respiratory Society (ATS/ERS) standards and recommendations (47), and percent predicted spirometric parameters were calculated using Global Lung Function Initiative (GLI) spirometry race-neutral reference equations (48). FENO was measured in accordance with ATS guidelines (49). History of a severe asthma exacerbation was defined as an increase in asthma symptoms requiring: (a) systemic corticosteroids for ≥ 3 consecutive days, or (b) hospitalization or emergency department visit for asthma symptoms (11).

### Human rhinovirus infection

BEC cultures were infected on the apical surface with human rhinovirus A16 (RV-A16, Source: ATCC®) at a multiplicity of infection (MOI) of 0.5. Viral suspension was removed from the apical surface after 2 hours of incubation with no subsequent rinsing. Viral load was measured in BECs at 2 days(d), 4d, 7d, and 10d post infection using Genesig® Human Rhinovirus Subtype 16 PCR Kit (Primerdesign®).

### Scanning electron microscopy

Transwells were fixed in 2% paraformaldehyde and 2% glutaraldehyde in 0.1M phosphate buffer at 4C°, then rinsed with 0.1M phosphate and post-fixed in 1% osmium tetroxide overnight at 4C°. Following three 10-minute washes with water, samples were gradually dehydrated to 100% EtOH at 10% increments. A biopsy punch tool was used to remove the transwell membranes to be critical point dried. Samples were then gold sputter-coated and imaged using a FEI Quanta 200F scanning electron microscope.

### Immunofluorescence microscopy

AECs cultured in transwells were fixed in 10% neutral buffered formalin (NBF) and washed in PBS before being embedded in paraffin. After sectioning and deparaffinization, samples were permeabilized with 0.3% Triton-X 100 for 5 minutes, treated with 20 µg/mL proteinase K (Invitrogen #25530049) for 6 minutes for antigen retrieval, and blocked with 10% normal goat serum (NGS Vector Laboratories, #S-1000-20) for 1 hour at room temperature (RT). Sections were incubated with rabbit anti-e-cadherin (Cell Signaling Technology #3195, 1:50) and mouse anti-Tubulin A4A (Invitrogen #PA5-105102,1:200) in staining buffer (PBS+1% NGS + 0.1% Triton-X 100) overnight at 4°C, washed 3 times in PBS-T (PBS with 0.1% Tween-20), and incubated with Alexa Fluor 488-conjugated goat anti rabbit (Invitrogen #A32731, 1:1000) and Alexa Fluor 594-conjugated donkey anti mouse (Invitrogen #A-21203, 1:1000) antibodies. After washing, sections were re-blocked in 10% NGS and incubated with mouse anti-dsRNA (SCICONS, clone J2; used at 1:100) pre-conjugated to Alexa Fluor 647 with Zenon^TM^ mouse IgG_2a_ labeling kit according to the manufacturer’s instructions (Invitrogen #Z25108). Stained sections were mounted with ProLong^TM^ Gold with DAPI (Invitrogen, #P36935) and allowed to cure overnight. Slides were then imaged on a Leica DM6000 widefield microscope and images were processed using ImageJ.

### Protein analysis

Protein concentrations of CXCL10, IFN-β, IFN-λ2/IL-28A, and IFN-λ3/IL-28B were measured in supernatant from uninfected BEC cultures and BEC cultures 2-days following RV infection via a Human Luminex® Discovery Assay (R&D®). IFN-β concentrations were found to be below the assay detection limit in all samples from uninfected BECs and in many samples following RV infection (data not shown). IFN-λ2 and IFN-λ3 were also below detection limits in many uninfected samples but were consistently detected following RV infection. For analyte concentrations above the assay detection limit in neat samples following RV infection, supernatant was assayed following a 10-fold dilution in Luminex® assay diluent.

### RNA collection

RNA was collected from BEC cultures pre-infection and at 2d, 4d, 7d, and 10d post RV infection. To collect RNA from BEC cultures, media was first removed from the basolateral chamber of transwells. 800 µL of lysis buffer (Invitrogen®, Product No. AM1912) was added to the apical surface of cultures. A pipet tip was then used to gently scratch each apical well in a crosshatch pattern to loosen BECs from the transwell membrane. RNA was extracted using the RNAqueous™ kit for total RNA isolation (Thermo Fisher Scientific).

### Interferon-β and CXCL10 treatment experiments

BEC cultures from SE (n=6) and NSE (n=6) donors were treated with IFN-β or CXCL10 starting 2 days prior to RV infection and continued for 2 days following RV infection. Recombinant human recombinant human IFN-β (1 ng/mL; Abcam^Ⓡ^) or CXCL10 (2 ng/mL; R&D Systems^Ⓡ^) was added to BEC ALI medium starting 2 days prior to RV infection and refreshed with each medium change (every other day). Two days following initiation of treatment with IFN-β or CXCL10 BEC cultures were infected with human RV-A16 at an MOI of 0.5 as described above. RNA was collected from BEC cultures pre-infection and 2d following RV infection.

### RNA sequencing and data processing

Total RNA was used to construct libraries by using the SMART-Seq v4 Ultra Low Input RNA Kit for Sequencing (Takara, San Jose, Calif), with reverse transcription followed by PCR amplification to generate full-length amplified cDNA. Sequencing libraries were constructed by using the NexteraXT DNA sample preparation kit with unique dual indexes (Illumina, San Diego, Calif) to generate Illumina-compatible barcoded libraries. Libraries were pooled and quantified using a Qubit Fluorometer (Thermo Fisher Scientific, Waltham, Mass). Sequencing of pooled libraries was carried out on a NextSeq 2000 sequencer (Illumina) with paired-end 53-base reads with a target depth of 5 million reads per sample. Base calls were processed to FASTQs on BaseSpace (Illumina), and a base call quality-trimming step was applied to remove low-confidence base calls from the ends of reads. Resulting bcl files were deconvoluted and converted to fastq format using Casava from Illumina. Fastq files were aligned to the Ensembl version of the human genome (GRCh38, Ensembl 91) by using STAR (version 2.4.2a). HTSeq-count (version 0.4.1) was used to generate gene counts with mode as “intersection (nonempty)” and minimum alignment quality set to 20 and otherwise set to default parameters. Quality metrics were compiled from PICARD (version 1.134), FASTQC (version 0.11.3), Samtools (version 1.2), and HTSeq-count (version 0.4.1). For quality control, samples that had human aligned counts greater than 1 million mapped reads and a median coefficient of variation coverage less than 0.7 were kept. Genes were filtered to include those that had a trimmed mean of M value normalization count of at least 0.3 in at least 10% of samples and further filtered for only protein coding genes. Normalized counts were transformed to log2 counts per million mapped reads along with observations level weights by using voomWithQualityWeights from the limma R package (version 4.2.1). The final data set for the time series analysis of SE vs NSE vs HC donors included 198 samples composed of 16275 genes and for the Interferon-β and CXCL10 conditioning experiments 48 samples composed of 15907 genes.

### Statistical analysis

To compare viral load differences over time among the SE, NSE, and HC groups a generalized additive mixed model (GAMM) was run with an interaction between time and group and a random effect for epithelial cell donorID. This was done using the R package “mgcv” with model syntax of log10(viral load) ∼ group + s(time) + s(time, by=group). The differences in means across time, abbreviated throughout as the Average Difference (Avg) are presented, as are the differences in the smoothing terms among groups (Shape). Differences in viral load restricted to the 2d timepoint were assessed using a linear model and the differences in means (Estimate) are presented.

To compare baseline protein levels between SE, NSE and Healthy, a linear model was used with model syntax: log10(protein analyte concentration) ∼ group. Association between baseline BEC CXCL10 protein concentration and post infection viral load was analyzed using a linear model with syntax: log10(viral load) ∼ log10(CXCL10 protein analyte concentration) with epithelial cell donorID as a random effect; cross sectional association to day 2 viral load was analyzed using Pearson correlation. GSEA analysis was performed using the Hallmark gene sets from the Molecular Signatures Database (MSigDB). Genes were ranked based on log2 fold change for the two exacerbation groups (SE and NSE) at baseline. GSEA was performed using the fgsea package in R with 10,000 gene set permutations to assess statistical significance. Enrichment scores (NES) and adjusted p-values were reported, and significant pathways were identified using a false discovery rate (FDR) cutoff of 0.2 or below by the Benjamini-Hochberg procedure.

Differentially expressed genes were identified using generalized additive mixed models (GAMMs) comparing expression differences over time between the two exacerbation groups (SE and NSE) as an interaction term on the first half (99 of 198) samples. The model syntax was: gene_expression ∼ exacerbation_group + s(time) + s(time,by=exacerbation_group) and including a random effect for epithelial cell donorID. This identified 6,714 genes that reached a false discovery rate adjusted p-value (FDR) < 0.25 by the Benjamini-Hochberg procedure for any of the 3 fixed effect terms. These genes were then utilized for supervised weighted gene co-expression network analysis (WGCNA) to identify genes with similar expression patterns and group them into modules. The WGCNA parameters used were minimum module size of 20, maximum module size of 500, deep split of 3, and a soft threshold (power) of 10, which was selected as the lowest soft threshold resulting in a scale free topology model fit R-squared >0.8. This resulted in 42 modules for downstream analysis. Module values were summarized by taking the mean of all genes (log2 transformed values) in the respective module. Using the gene composition of each module these modules were generated for all samples for all the downstream analysis. These modules were then modeled using the same GAMM syntax to identify those showing differential expression patterns over time among SE, NSE, and HC groups at a stringent FDR < 0.05 for the interaction and/or group intercept terms. Multiple testing correction was performed using the Benjamini-Hochberg procedure. For each module that showed a significant difference among groups, pathway enrichment was performed using the clusterprofiler R package, which calculates a hypergeometric FDR corrected p-value for enrichment of public genesets. We used the gene ontology biological processes (GO_BP), KEGG, Reactome, Biocarta, and MSigDB Hallmark genesets for enrichment. The five modules presented in the main text were annotated based on manual inspection of the enrichment terms and module genes. Linear mixed effects models were used to identify modules significantly associated with viral load in the post infection samples using the R package “kimma”(50). The model syntax was: module_expression ∼ log10(viral load) + time and including a random effect for epithelial cell donorID. For the mediation analysis, 2 linear models were compared using only the post infection samples. The syntax of the first model was module_expression ∼ exacerbation + time and including a random effect for epithelial cell donorID to compare module expression between SE and NSE. The second model used the same syntax but with the addition of log10 transformed viral load as a covariate to regress out the effects of viral load. The effect size and p-values for each module were then compared between the two models. The above models were also run adjusting for sex where the results were congruent.

To determine the effect of IFN-β or CXCL10 on RV replication, viral load was quantified at 2 days post-infection and compared across 3 treatment conditions using a linear mixed model with syntax: log10(viral load) ∼ Treatment including a random effect for epithelial cell donorID separately for SE and NSE groups. A similar model was used to compare module expression across treatment conditions adjusted for median cv coverage using the syntax: module_expression ∼ Treatment + median_cv_coverage, including a random effect for donorID.

### Single cell analysis sample preparation

For scRNA-seq, BECs from a subset of the children with asthma (n=11) and healthy children (n=3) were differentiated ex vivo at ALI to generate organotypic cultures and infected with human rhinovirus A16 at an MOI of 0.5 as described above. To dissociate differentiated BEC ALI cultures into a single cell suspension, the apical surface of the transwell was treated with 0.5mL of 10mM dithiothreitol (DTT; Thermo Scientific™) in 1xPBS (Gibco) for 10 minutes at 37°C to remove accumulated mucus, subsequently washed three times with 0.5mL of 1xPBS (Gibco) to remove excess DTT, and then treated with 300uL TrypLE Express (Gibco) warmed to 37°C, with 600uL TrypLE Express added to the basolateral chamber. Wells were incubated for 15 minutes at 37°C, then mechanically dissociated by pipetting 10 times with a P1000 pipet. The cell suspension was transferred into an empty 15mL conical tube, where the cell suspension was further pipetted until visually homogeneous, and wells were rinsed with warm EX Plus media (StemCell). Cells were mixed by pipetting 20 times with a 5mL serological pipet set to slow and then centrifuged at 250x g for 7 mins. The cell pellet was resuspended with 1mL of 0.04% BSA (Invitrogen) in 1xPBS and gently pipetted 30 times and an additional 1mL of 0.04% BSA in PBS was added. The cell suspension was then filtered twice through a 30-µm cell strainer (pluriSelect) and centrifuged at 150x g for 10 minutes. The 10X Genomics Chromium Next GEM Single Cell Fixed RNA Sample Preparation Kit was used to fix the BECs. The cell pellet was resuspended in 1mL of Fixation Buffer (10X Genomics), pipetted to mix 10 times, and stored overnight at 4°C. The fixed single cells were then centrifuged 850x g for 7 mins., resuspended with 1mL chilled Quenching Buffer (10X Genomics), and pipetted to mix 5 times on ice. The cell suspension was transferred to 2mL lo-bind tubes (Eppendorf), 100uL Enhancer (10X Genomics) warmed to 65°C was added and mixed, and 265uL 50% glycerol was added and mixed into the fixed cells. Samples were stored at -80°C until further post-storage processing and sequencing.

Fixed samples were processed according to manufacturer’s instructions, using the Chromium Next GEM Single Cell Fixed RNA Sample Preparation Kit (10x Genomics). Briefly, fixed samples were thawed and hybridized overnight to one of four human probe sets (10x Genomics), with or without additional spiked-in custom rhinovirus probes. Pooled samples were washed and loaded onto the Chromium X (10x Genomics) for partitioning into GEMs. Gene expression libraries were generated according to the Chromium Fixed RNA Profiling Reagent Kits for Multiplexed Samples user guide. Sequencing was carried out on a NextSeq 2000 sequencer, using NextSeq 2000 P4 XLEAP-SBS flowcells (Illumina) with a target depth of 10,000 reads/cell.

Custom RV probes for 10x Genomics Flex were designed targeting the human RV A16 genome (GenBank: L24917.1). Following 10x specifications, probes consisted of 50 bp target sequences split into 25 bp left (LHS) and right (RHS) segments, with 44–72% GC content, a T at the 3′ end of the LHS, no homopolymer runs >3, and minimal target site competition. Primer-BLAST was used to identify candidate sequences with human genome specificity (taxID:9606). From 40 valid targets, eight met all criteria including a 25th-position T; three were excluded due to binding overlap, leaving five. Reverse strand primers were reverse-complemented to generate RHS probes. Final sequences were manually verified for human genome specificity. A sixth validated probe was added based on extensive use in RV qPCR work (51). Probe sequences are shown here:

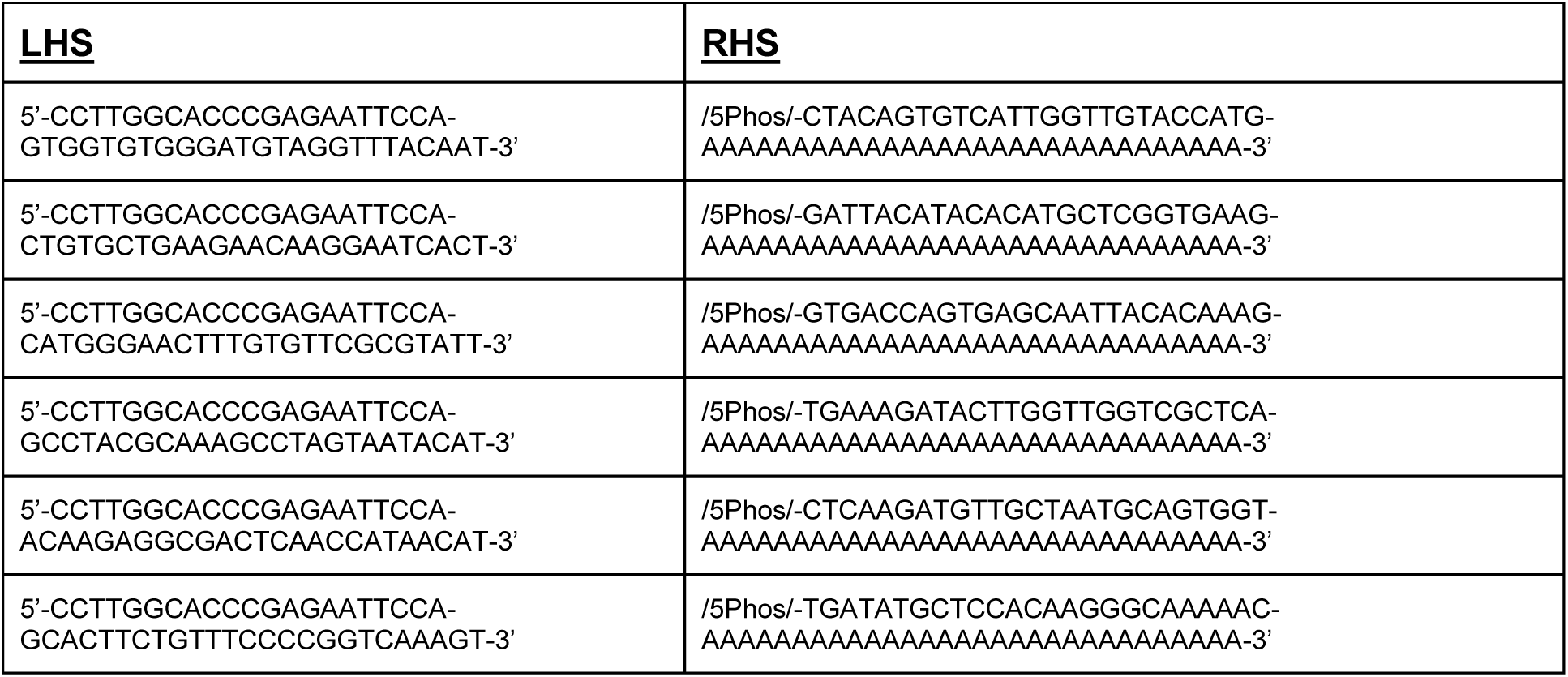

### Single cell RNA-seq data processing and analysis

Single-cell RNA-sequencing (scRNA-seq) 10x dataset preprocessing including cell demultiplexing and alignment were performed using Cell Ranger Single-Cell Software Suite (version 7.0.0, 10x Genomics Inc). We used the human reference genome (GRCh38, Ensembl 91) for alignment and generated raw and filtered cell by gene count matrix files.

The filtered cell-by-gene count matrix file was used for the standard Cell Ranger based data processing pipeline. First, we removed genes expressed in fewer than four cells. Next, we excluded cells with fewer than 100 genes detected or cells with more than 20% mitochondrial reads. Further, to eliminate possible doublet cells, we removed cells with more than 7000 genes detected. The counts data were then normalized to 10,000 reads per cell using the NormalizeData function. Samples were integrated using a stepwise canonical correlation analysis (CCA) approach with the IntegrateLayers function on 20 dimensions and 2,000 variable genes. After integration, principal component analysis (PCA) was performed on the integrated data using the RunPCA function with the first 50 principal components. This was followed by Uniform Manifold Approximation and Projection (UMAP) using RunUMAP function with 50 dimensions, and clustering was performed by using the FindNeighbors and FindClusters functions with a resolution of 0.85 to identify distinct cell clusters. Cluster specific markers for each cluster were identified using FindAllMarkers function, then cell clusters were manually annotated by assessing the expression of cluster specific markers to known cell type markers. Mitotic basal cells were annotated by the expression of cell cycle markers such as TOP2A, MKI67, and NUSAP1; proliferative basal cells were annotated by MCM4, MCM5, and HELLS; basal cells by the expression of TP63, KRT5, KRT15, and BCAM; and suprabasal cells by NOTCH3 and KRT19. We identified secretory cells by unique expression of SERPINB3 and CLCA2, and goblet cells by MUC5AC, MUC5B and VMO1. We identified three cell clusters within ciliated cells, deuterosomal cells by CCNO, DEUP1, and CDC20B, and ciliated cells by FOXJ1, PIFO, CDHR3, SNTN and DNAH9, and mucous-ciliated cells expressed a combination of ciliated markers and lower levels of mucin genes such as MUC5AC and MUC5B. We also identified rare cell populations such as ionocytes by the expression of FOXI1, CFTR and ASCL3, and tuft (brush) cells and PNECs by the expression of POU2F3, ASCL2 and CHGA and ASCL1. Additionally, we observed a distinct cluster representing cells in an inflammatory state with the expression of genes such as ISG20, IFIH1, IFI30, IFIT1, IFIT2, IFIT3 and CXCL10. All the analyses were performed in the Seurat R package(52) (version 5.0.1).

### Study approval

BECs from children were obtained and utilized in these experiments under studies #12490 and #1596 approved by the Seattle Children’s Hospital Institutional Review Board. Parents of subjects provided written consent and children over 7 years of age provided assent.

## Data availability

All raw bulk and single cell RNA-sequencing data have been uploaded to the NCBI Gene Expression Omnibus database and accessible via *GSE309705*.

## Author contributions

JSD and MCA designed the studies. MPW recruited and enrolled human subjects. PCDC, LMR, ERV, CRG, NBS, KW, and JSD performed the experiments. NDJ, BB, PCDC, AJN, and MCA performed the RNA-sequencing analysis. PCDC, BB, WTP, NDJ, MCA, and JSD wrote the manuscript. PCDC, BB, WTP, NDJ, LMR, NBS, KW, AJN, ERV, CRG, MPW, TSH, SFZ, MCA, and JSD edited the manuscript. All authors reviewed and approved the manuscript prior to submission.

## Acknowledgements

Grant Funding Sources: The work presented in this manuscript was supported by the following National Institutes of Health (NIH) grants: NIH R01AI163160 (JSD), NIH K24AI150991(JSD), NIH U19AI175089 (TSH, JSD, MCA).

We would like to acknowledge the BRI Genomics Core, RRID:SCR_026658 for use of their facilities for generating and processing of NGS data.

## Supplemental material

**Supplemental Figure 1.**
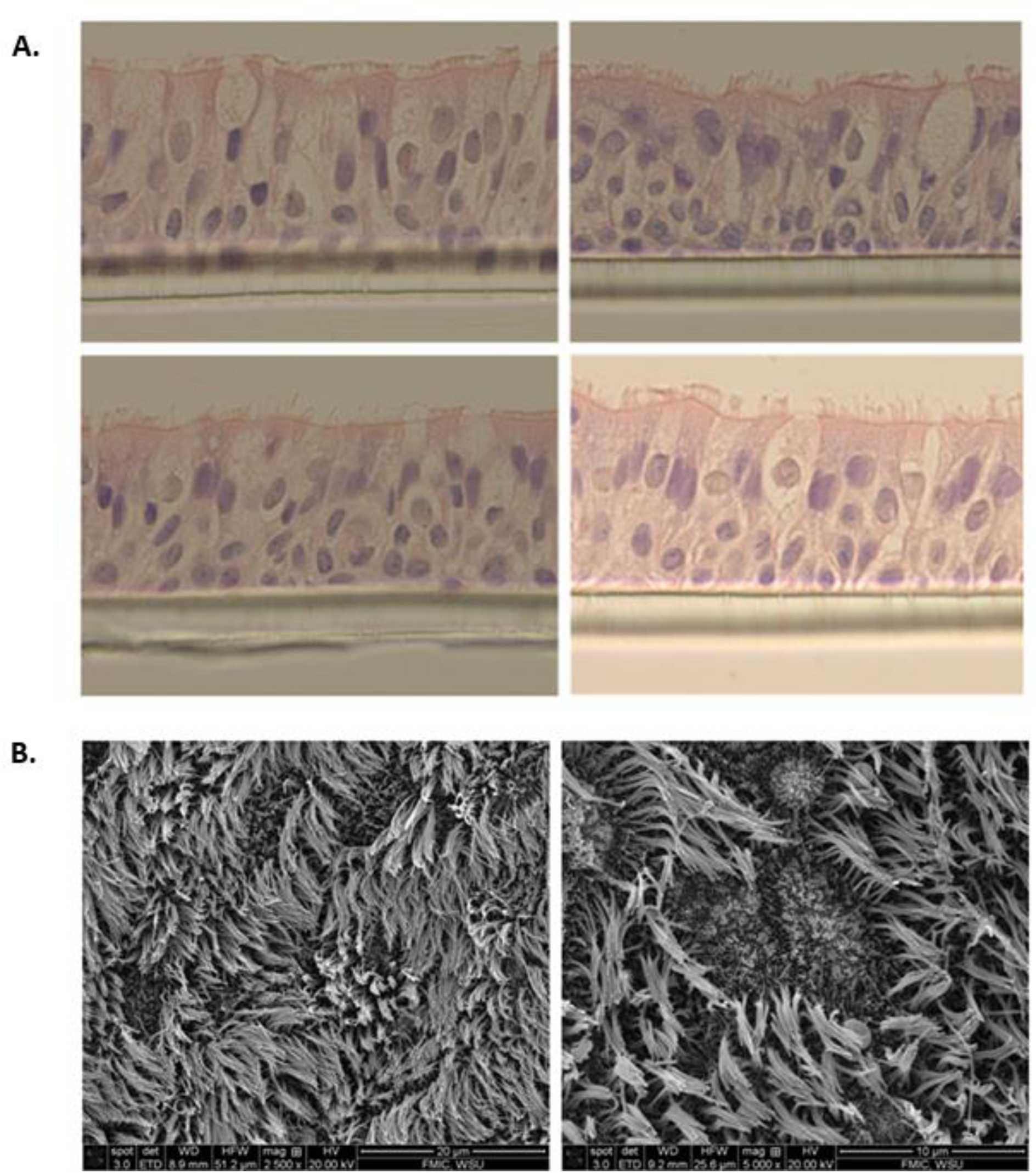
Organotypic bronchial epithelial cultures. (**A**) Representative hematoxylin and eosin (H&E) stained histology cross sections and (**B**) scanning electron microscopy images of passage 3 primary bronchial epithelial cultures from study asthma donors differentiated to an organotypic pseudostratified and ciliated state for 21 days under air-liquid interface conditions using PneumaCult™ALI medium (Stemcell™).

**Supplemental Figure 2.**
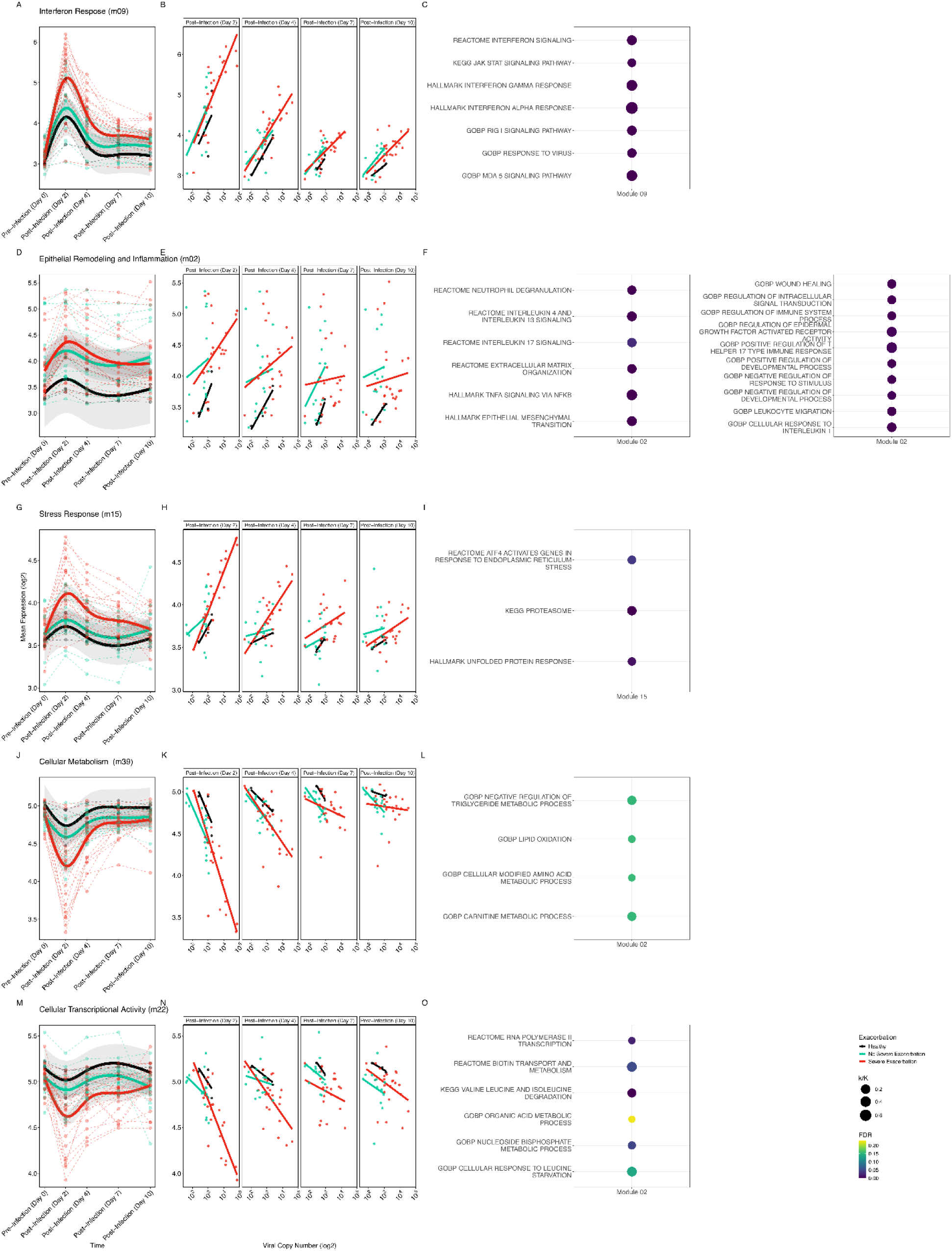
Time course of Module Expression and Correlation with Viral Load, with Functional Enrichment Annotations. (**A, B**) Scatterplots showing the non-linear changes in expression of the “Interferon Response” module over time by donor group. BECs from SE Asthma donors exhibited a significantly higher average expression level at 2 days, with a 1.69-fold increase compared to NSE Asthma donors and a 1.92-fold increase compared to Healthy donors with a significantly different GAMM shape fit characterized by sustained higher levels at each subsequent timepoint (SE vs NSE: GAMM All timepoint: FDR=3.48e-05, linear model effect size restricted to d2: Estimate=0.76, FDR=9.47e-03, SE vs Healthy: GAMM All timepoint: FDR=9.87e-07, linear model effect size restricted to d2: Estimate=0.94, FDR=1.35e-1). (**C**) Selected enriched GO, Biocarta, Reactome, MSigDB Hallmark and KEGG terms for the 363 genes within the “Interferon Response’’ module. Dot size denotes the k/K value (ratio of number of genes in module (k) divided by the number of genes in the indicated geneset (K)), and the color denotes statistical significance of the pathway enrichment at FDR ranging from 0-0.20. (**D, E**) Analogous plots of the “Epithelial Remodeling and Inflammation” module. BECs from SE Asthma donors demonstrated a 1.15-fold higher expression level at 2d compared to NSE donors and sustained elevation at 4d with return to baseline levels in the NSE asthma group (SE vs NSE: GAMM All timepoint: FDR=8.4e-03, linear model effect size restricted to d2: Estimate=0.20, FDR=6.50e-01), and SE Asthma donors demonstrated a sustained greater expression compared to Healthy donors at each timepoint with a 1.67-fold higher expression at 2d (SE vs Healthy: GAMM All timepoint: FDR=6.55e-03, linear model effect size restricted to d2: Estimate=0.74, FDR=2.87e-01). (**F, G**) Selected enriched GO, Biocarta, Reactome, MSigDB Hallmark and KEGG terms for the 1396 genes within the “Epithelial Remodeling and Inflammation’’ module. Dot size denotes the k/K value (ratio of number of genes in module (k) divided by the number of genes in the indicated geneset (K)), and the color denotes statistical significance of the pathway enrichment at FDR ranging from 0-0.20. (**H, I**) Analogous plots of the “Stress Response” module. BECs from SE Asthma donors demonstrated a 1.23-fold higher expression level at 2d compared to NSE donors and a 1.33-fold higher expression compared to Healthy donors, with sustained higher levels over time (SE vs NSE: GAMM All timepoint: FDR=1.8e-3, linear model effect size restricted to d2: Estimate=0.30, FDR=1.66e-02, SE vs Healthy: GAMM All timepoint: FDR=7.73e-04, linear model effect size restricted to d2: Estimate=0.41, FDR=1.53e-01). (**J**) Selected enriched GO, Biocarta, Reactome, MSigDB Hallmark and KEGG terms for the 242 genes within the “Stress Response’’ module. Dot size denotes the k/K value (ratio of number of genes in module (k) divided by the number of genes in the indicated geneset (K)), and the color denotes statistical significance of the pathway enrichment at FDR ranging from 0-0.20. (**K, L**) Scatterplot showing the non-linear changes in expression of the “Cellular Metabolism” module over time differing by clinical exacerbation group. BECs from SE Asthma donors demonstrated a significantly lower average expression level at 2 days, with a 1.30-fold decrease compared to NSE Asthma donors and a 1.52-fold decrease compared to Healthy donors with a significantly different GAMM shape fit characterized by sustained lower levels at each subsequent timepoint (SE vs NSE: GAMM All timepoint:FDR=8.1e-04, linear model effect size restricted to d2: Estimate=-0.38, FDR=1.66e-02, SE vs Healthy: GAMM All timepoint: FDR=9.80e-05, linear model effect size restricted to d2: Estimate=-0.60, FDR=1.12e-01). (**M**) Selected enriched GO, Biocarta, Reactome, MSigDB Hallmark and KEGG terms for the 106 genes within the “Cellular Metabolism’’ module. Dot size denotes the k/K value (ratio of number of genes in module (k) divided by the number of genes in the indicated geneset (K)), and the color denotes statistical significance of the pathway enrichment at FDR ranging from 0-0.20. (**N, O**) Analogous plots of the “Cellular Transcriptional Activity” module. BECs from SE donors demonstrated a significantly lower average expression level at 2 days, with a 1.21-fold decrease compared to NSE donors and a 1.32-fold decrease compared to Healthy donors with a significantly different GAMM shape fit characterized by sustained lower levels at each subsequent timepoint (SE vs NSE: GAMM All timepoint:FDR =6.8e-04, linear model effect size restricted to d2: Estimate=-0.28, FDR=1.66e-02, SE vs Healthy: GAMM All timepoint: FDR=2.40e-03, linear model effect size restricted to d2: Estimate=-0.41, FDR=1.35e-01). (**P**) Selected enriched GO, Biocarta, Reactome, MSigDB Hallmark and KEGG terms for the 204 genes within the “Cellular Transcriptional Activity’’ module. Dot size denotes the k/K value (ratio of number of genes in module (k) divided by the number of genes in the indicated geneset (K)), and the color denotes statistical significance of the pathway enrichment at FDR ranging from 0-0.20.

**Supplemental Figure 3.**
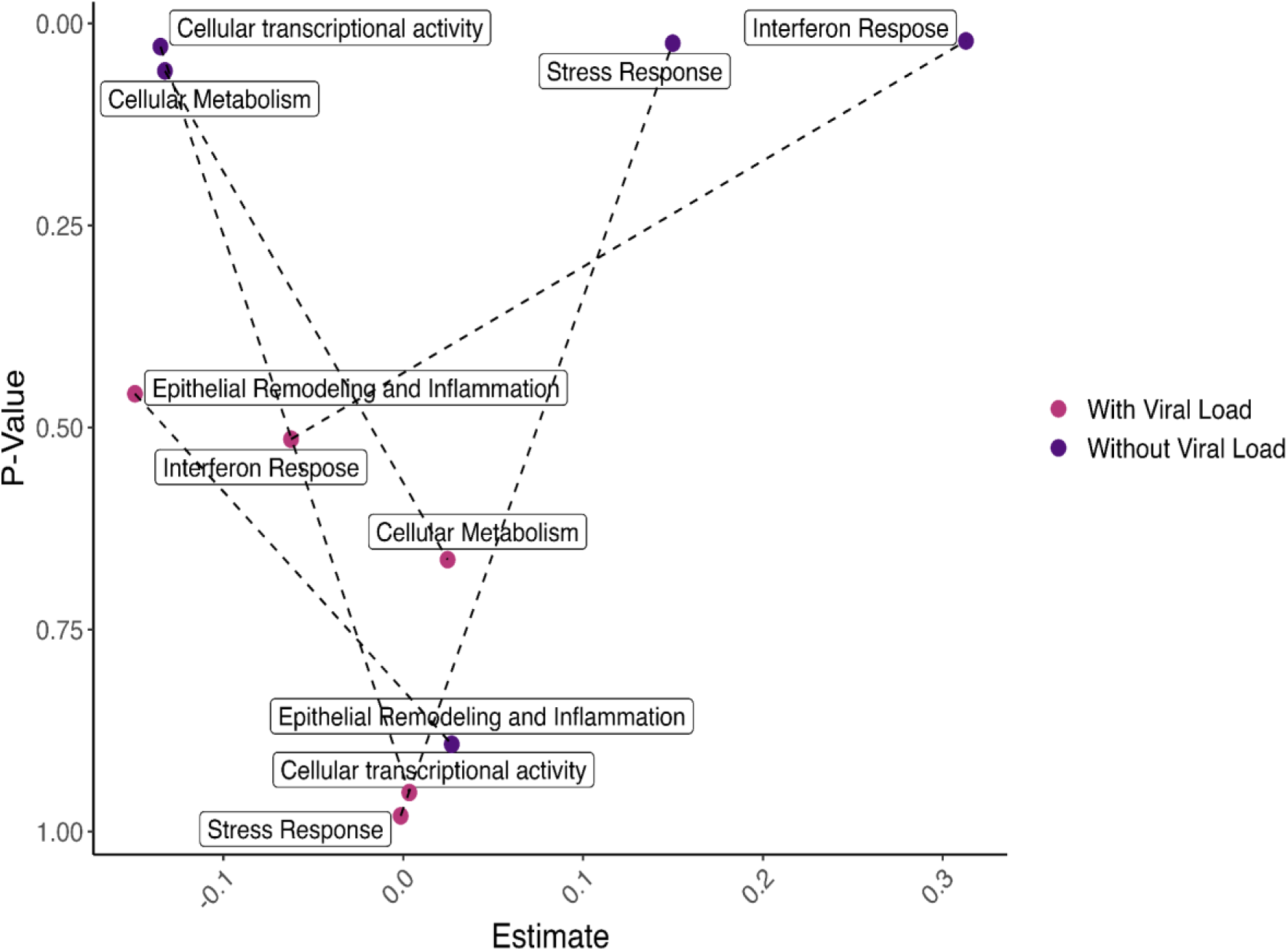
Viral load has a mediating effect on Module expression. A linear model subset to the post infection timepoints, d2-10, was used to assess the extent to which viral load differences were mediating the magnitude differences in module expression observed between SE and NSE groups. Four of the modules of interest, Interferon response, Stress response, Cellular metabolism, and Cellular transcriptional activity, showed that viral load mediated the majority of difference in module expression observed between the two groups. The differences between SE and NSE in the Interferon response module were: without viral load: Estimate=0.42, p=9.56e-03; with viral load: Estimate=-0.045, p=0.69. Stress response module: without viral load: Estimate=0.14, p=0.024; with viral load: Estimate= -0.0014, p=0.98. Cellular metabolism module: without viral load: Estimate=-0.13, p=0.058; with viral load: Estimate=0.024, p=0.66. Cellular transcriptional activity module: without viral load: estimate=-0.13, p=0.028; with viral load: Estimate=0.0033, p=0.95. No result could be derived for the Epithelial Remodeling and Inflammation module as this module largely differed by kinetic shape between groups rather than overall magnitude of expression Shown are the model estimates on x-axis and p-values on the y-axis for the model without (purple) or with (pink) viral load included in the model. The 5 modules of interest are labeled.

**Supplemental Figure 4.**
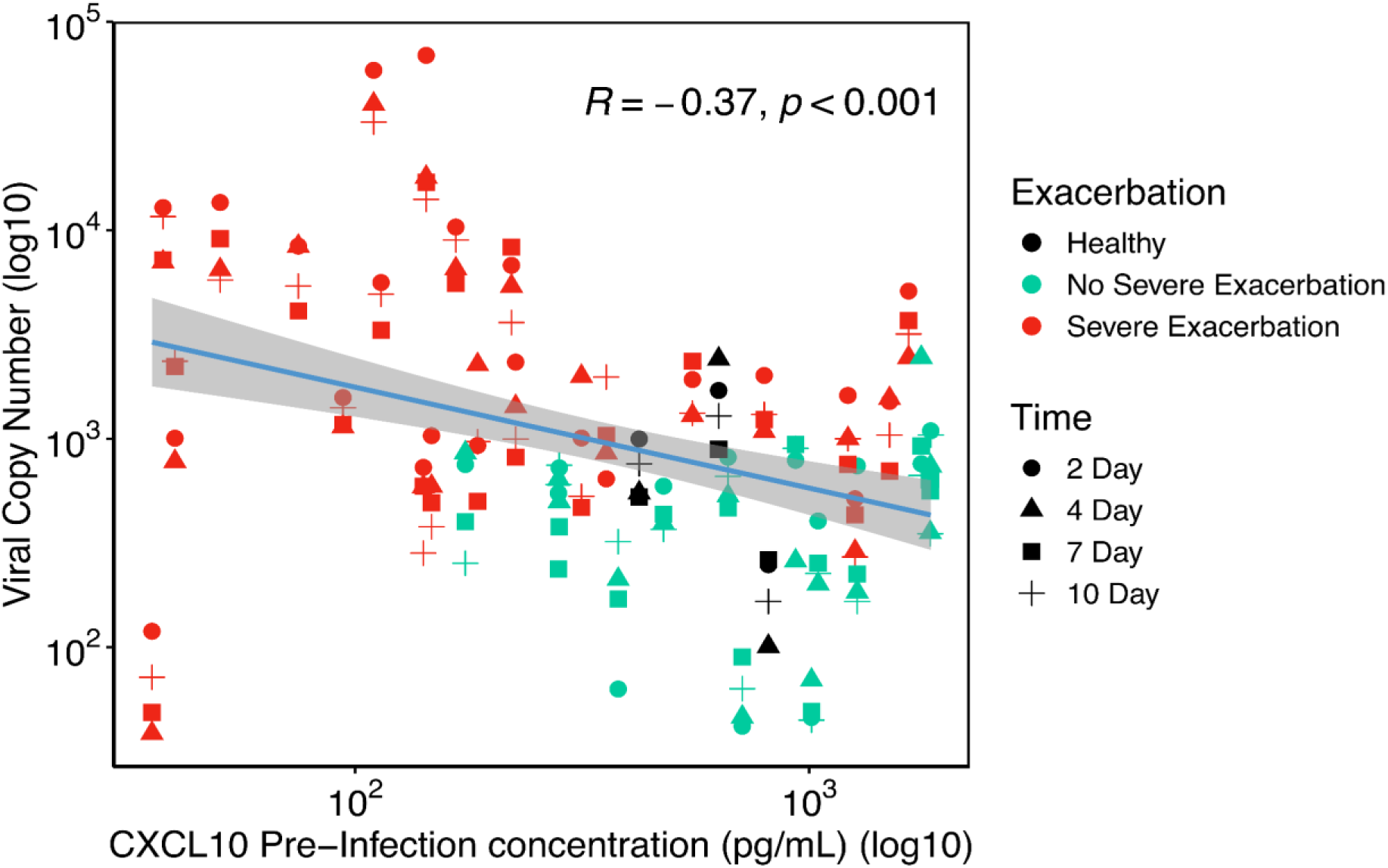
Post-Infection Viral load shows a significant inverse relationship to Pre-Infection CXCL10. Scatter plot showing a significant inverse relationship between log transformed pre-infection CXCL10 protein secretion and log transformed viral load post-infection d2-d10.

**Supplemental Figure 5.**
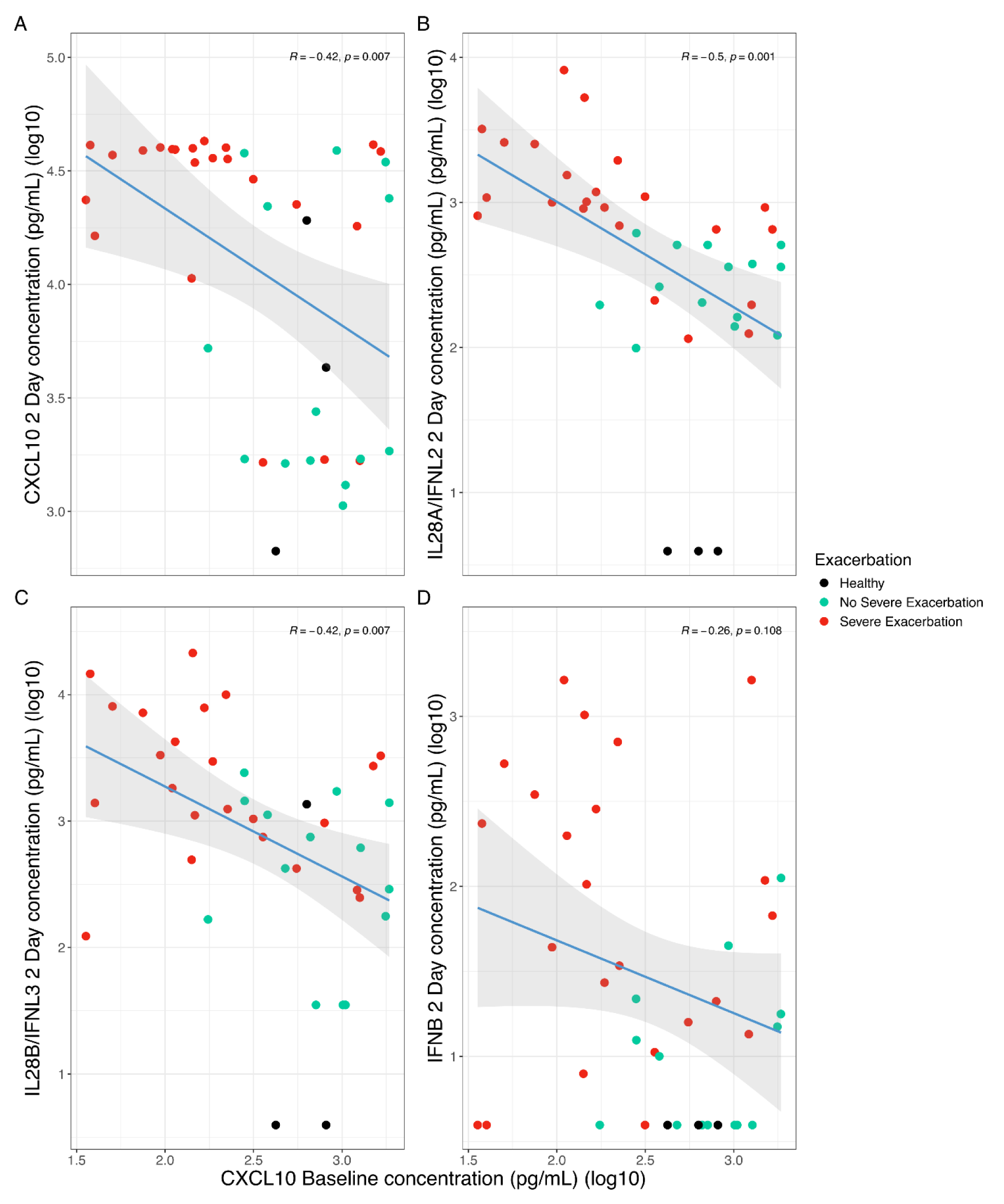
Pre-Infection CXCL10 shows a significant inverse relationship to Day 2 CXCL10, IFNβ, IFN-λ2/IL-28A and IFN-λ3/IL-28B. Scatter plot showing a significant inverse relationship between log transformed pre-infection CXCL10 protein secretion and subsequent protein secretion at 2 days post RV infection of (**A**) CXCL10 (Estimate=-0.37, p=7.16e-03), (**B**) IFN-λ2 (Estimate=-0.69, p=2.32e-05), and, (**C**) IFN-λ3 (Estimate=-0.34, p=4.24e-03) as well as a non-significant inverse trend with (**D**) IFN-β (Estimate=-0.14, p=1.69e-01). Fit lines are based on a linear model including 95% confidence intervals.

**Supplemental Figure 6.**
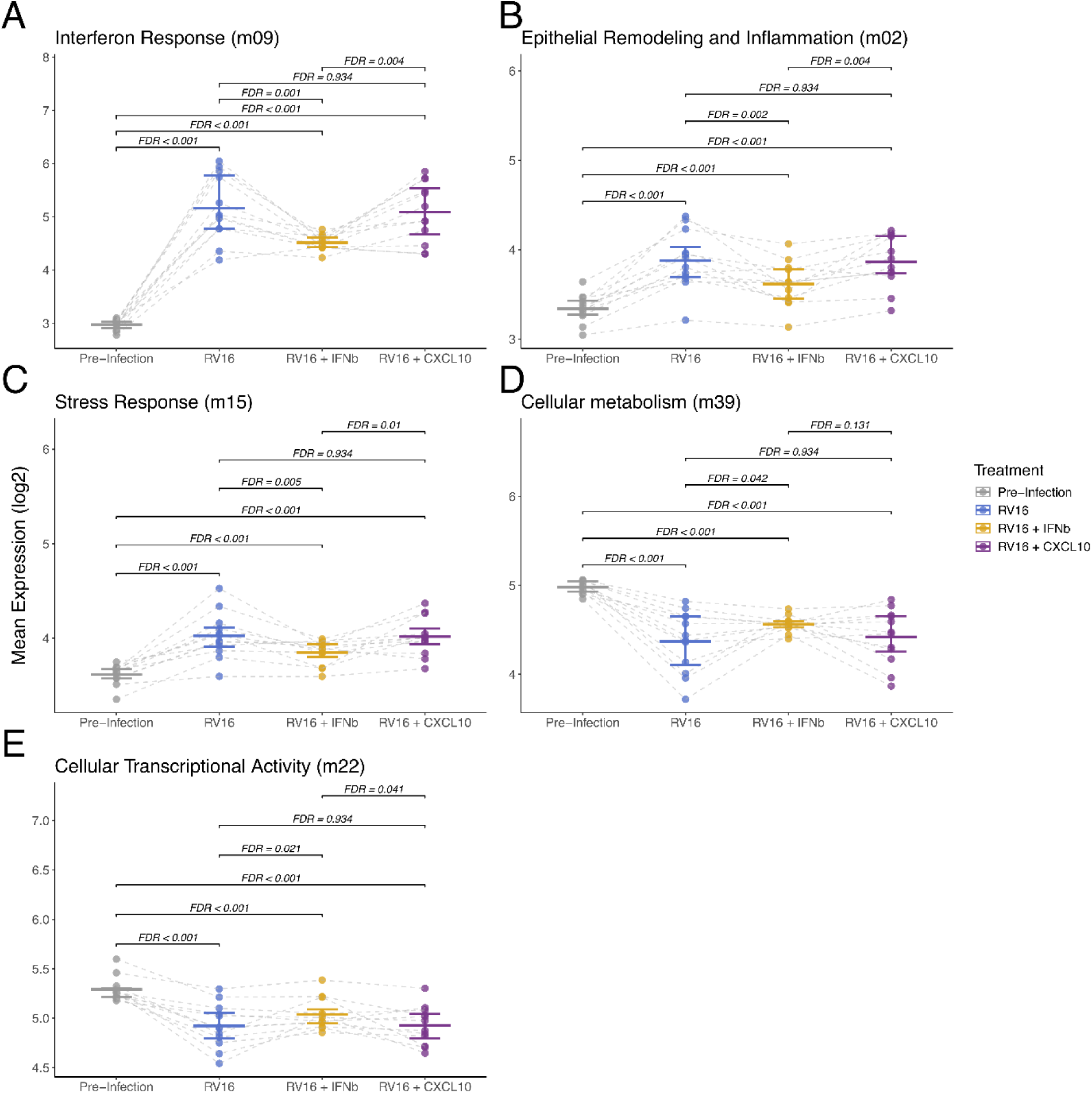
Module Expression Difference by RV16, RV16 + IFN-β and RV16 + CXCL10. (**A**) Dot plot showing expression changes in Interferon response module by IFN-β treatment showing a 1.53-fold decrease in expression with IFN-β treatment compared to RV16 (estimate=-0.6, p=1.23e-03) and whereas there is a 4.42-fold and a 2.88-fold increase in expression from Pre-infection compared to RV16 infected sample and IFN-β treated samples respectively (RV16:estimate=2.14, p=2.74e-15, IFN-β:estimate=1.52, p=4.25e-11). (**B**) Dot plot showing expression changes in Epithelial remodelling and Inflammation response module by IFN-β treatment showing a 1.20-fold decrease in expression with IFN-β treatment (estimate=-0.27, p=2.24e-03) and whereas there is a 1.47-fold and a 1.21-fold increase in expression from Pre-infection compared to RV16 infected sample and IFN-β treated samples respectively (RV16:estimate=0.55, p=9.41e-09, IFN-β:estimate=0.28, p=3.23e-04). (**C**) Dot plot showing expression changes in Stress response module by IFN-β treatment showing a 1.14-fold decrease in expression with IFN-β treatment (estimate=-0.19, p=4.61e-03) and whereas there is a 1.35-fold and a 1.18-fold increase in expression from Pre-infection compared to RV16 infected sample and IFN-β treated samples respectively (RV16:estimate=0.43, p=1.39e-08, IFN-β:estimate=0.23, p=2.41e-04). (**D**) Dot plot showing expression changes in Cellular metabolism module by IFN-β treatment showing a 1.12-fold increase in expression with IFN-β treatment (estimate=0.16, p=4.20e-02) and whereas there is a 1.48-fold and a 1.32-fold decrease in expression from Pre-infection compared to RV16 infected sample and IFN-β treated samples respectively (RV16:estimate=-0.56, p=1.18e-08, IFN-β:estimate=-0.40, p=5.64e-06). (**E**) Dot plot showing expression changes in Cellular transcriptional activity module by IFN-β treatment showing a 1.09-fold increase in expression with IFN-β treatment (estimate=0.13, p=2.13e-02) and whereas there is a 1.31-fold and a 1.19-fold decrease in expression from Pre-infection compared to RV16 infected sample and IFN-β treated samples respectively (RV16:estimate=-0.39, p=1.18e-08, IFN-β:estimate=-0.25, p=1.57e-05). Boxplots indicate median and interquartile ranges for each group and the Fit line is based on a linear model including 95% confidence intervals.

**Supplemental Figure 7.**
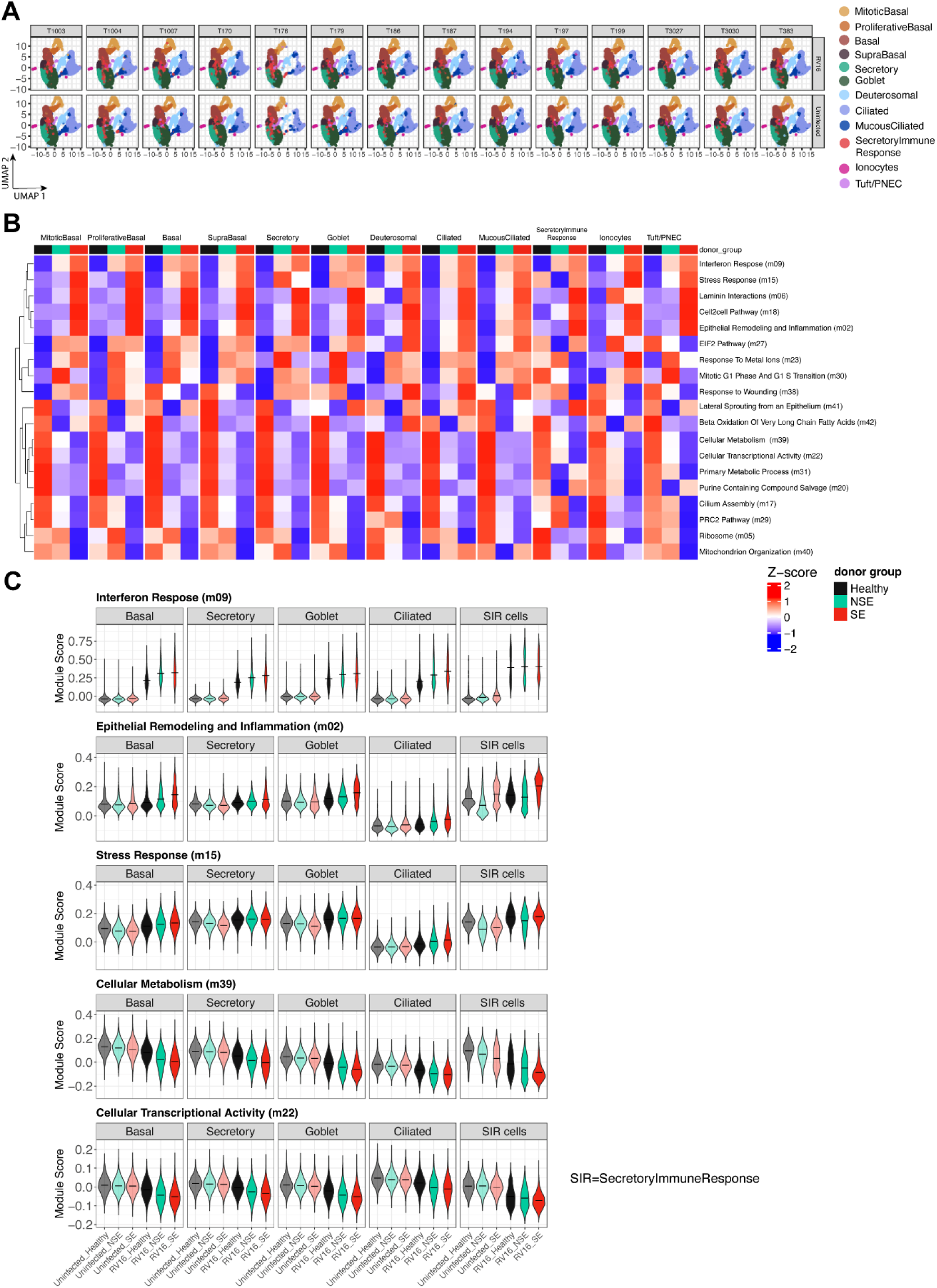
Single cell transcriptomics, module level analysis. (**A**) UMAP representation of the 316,712 cells separated by donor and condition (uninfected and RV infection). (**B**) Heatmap showing relative expression of modules (defined by WGCNA from bulk transcriptomics) expression at 48 hr following RV infection across donor groups, healthy, non-Severe and Severe at individual cell types. Module expression levels are shown as row normalized Z-scores of the mean expression for each group with red representing higher relative expression and blue representing lower relative expression. Module row ordering is by hierarchical clustering. (**C**) Violin plot showing distribution of module expression for select modules by condition (uninfected and RV infection) by donor groups, healthy, non-Severe and Severe at individual cell types. The black horizontal line in each violin denotes a mean expression.

**Supplemental Figure 8.**
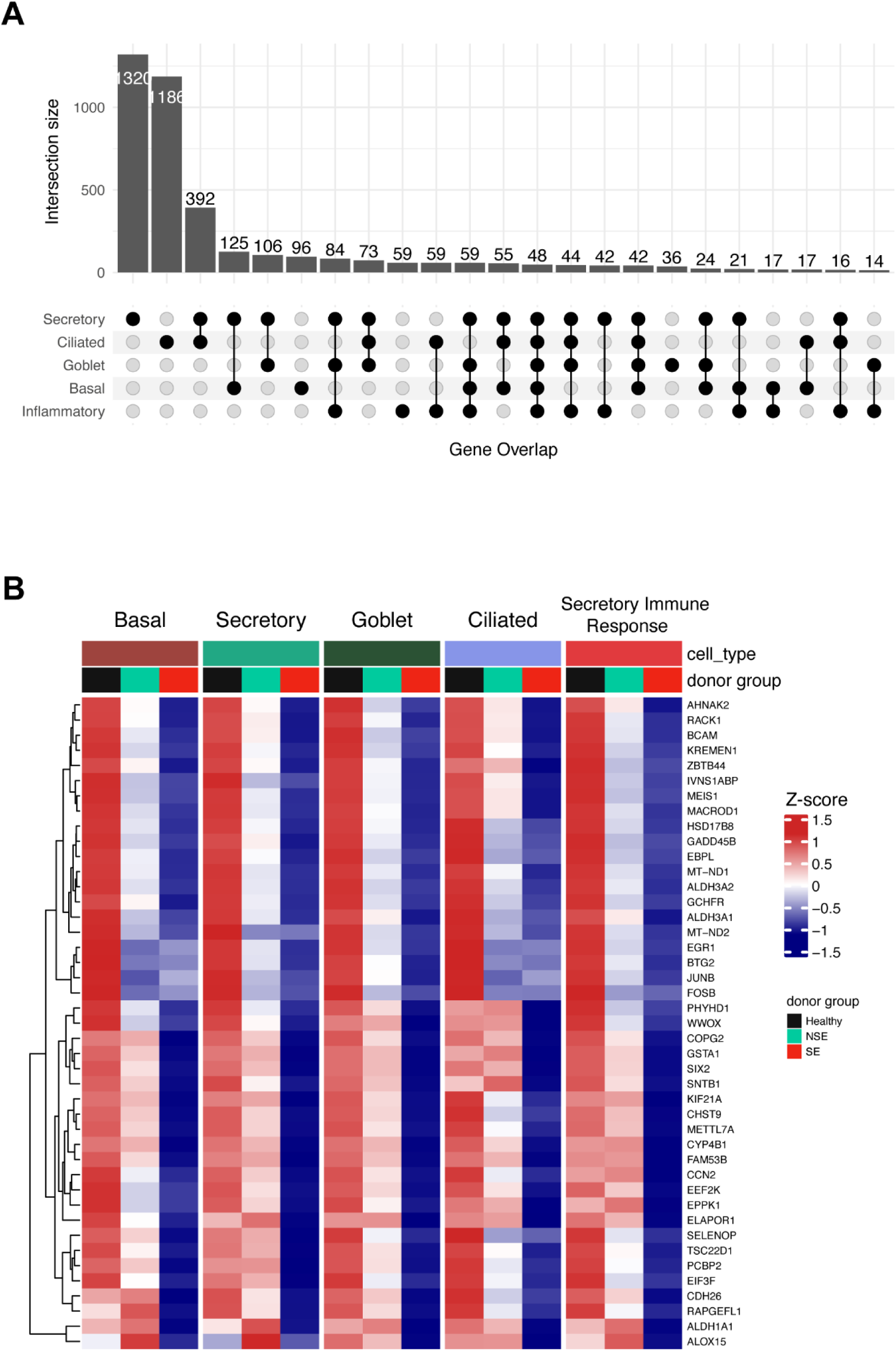
Single cell transcriptomics, gene level analysis. (**A**) Upset plot showing the count of overlapping genes with descending trend across five cell types, basal, secretory, goblet, ciliated and secretory immune response cells. (**B**) Heatmap showing relative expression of 48 core signature genes (shared across all five cell types) with descending expression trends at 5 cell types. Gene expression levels are shown as row normalized Z-scores of the mean expression for each group with red representing higher relative expression and blue representing lower relative expression.

## Supplemental Tables

**Supplemental Table 1.** Generalized Additive Mixed Models (GAMMs) results comparing expression differences over time between healthy and the two exacerbation groups. Table shows the smoothing terms of time and smoothing interaction term for the effect of exacerbation groups and healthy samples over time and shows the average difference by the exacerbation and healthy groups.

**Supplemental Table 2.** Results of the Linear Model shows the differential expressed modules by viral load over time and subset to Day 2

**Supplemental Table 3A:** Module annotations based on gene enrichment

**Supplemental Table 3B:** Table of genes in Modules EnsembleIDs

**Supplemental Table 3C:** Table of genes in Modules hgnc symbol

**Supplemental Table 4.** Linear Model results comparing expression at Day 2 between healthy and the two exacerbation groups.

**Supplemental Table 5.** Linear mixed effects models results to check for mediation effect of viral load

**Supplemental Table 6.** Linear model comparison of baseline BEC supernatant concentrations by exacerbation group

**Supplemental Table 7.** Linear Model results comparing expression difference between RV16, IFNb and CXCL10 Treatment groups.

**Supplemental Table 8.** Linear Model comparison of viral copy number to the module expression.

**Supplemental Table 9.** Linear mixed effects models results to check for mediation effect of viral load with Treatment group RV16, RV16 + IFNb and RV16 + CXCL10

**Supplemental Table 10. Cell Type-Specific Marker Genes.** This table lists the top marker genes that distinguish each cell type from the others, as identified through differential expression analysis. These markers were used to define and validate cell type annotations across the dataset.

**Supplemental Table 11. Cell Counts by Donor, Condition, and Cell Type.** This table provides a breakdown of cell counts stratified by individual donor ID, donor group (healthy controls, NSE asthma, SE asthma), experimental condition (uninfected vs.RV-infected), and annotated cell type.

**Supplemental Table 12. CLM Module-Level Estimates in RV-Infected Samples** Contains output from the Cumulative Linked Model (CLM) ordinal regression framework, reporting model estimates for all modules across all cell types within the RV-infected condition.

**Supplemental Table 13. CLM Gene-Level Estimates in RV-Infected Samples** Contains output from the Cumulative Linked Model (CLM) ordinal regression framework, reporting model estimates for genes across all cell types within the RV-infected condition.

**Supplemental Table 14. Enriched Biological Terms from CLM-Identified Genes** Lists significantly enriched biological pathways and gene ontology terms derived from the set of genes identified as significant in the CLM model under RV-infected conditions.

